# Integrative molecular profiling of autoreactive CD4 T cells in autoimmune hepatitis

**DOI:** 10.1101/2020.01.06.895938

**Authors:** Amédée Renand, Iñaki Cervera-Marzal, Laurine Gil, Chuang Dong, Erwan Kervagoret, Hélène Auble, Sarah Habes, Anais Cardon, Jean-Paul Judor, Jean-François Mosnier, Sophie Brouard, Jérôme Gournay, Pierre Milpied, Sophie Conchon

## Abstract

**Background & Aims:** In most autoimmune disorders, crosstalk of B cells and CD4 T cells results in the accumulation of autoantibodies. In autoimmune hepatitis (AIH), the presence of anti-Soluble Liver Antigen (SLA or SepSecs) autoantibodies is associated with significantly reduced overall survival, but the associated autoreactive CD4 T cells have not been characterized yet. Here we isolated and deeply characterized SLA-specific CD4 T cells in AIH patients.

**Methods:** We used brief *ex vivo* restimulation with overlapping SLA-derived peptides to isolate and phenotype circulating SLA-specific CD4 T cells, and integrative single-cell RNA-seq (scRNA-seq) to characterize their transcriptome and TCR repertoire in n=5 AIH patients. SLA-specific CD4 T cells were tracked in peripheral blood through TCR sequencing, to identify their phenotypic niche. We further characterized disease-associated peripheral blood T cells by high content flow cytometry in an additional cohort of n=46 AIH patients and n=18 non-alcoholic steatohepatitis (NASH) controls.

**Results:** Autoreactive SLA-specific CD4 T cells were only detected in patients with anti-SLA autoantibodies and had a memory PD-1^+^CXCR5^−^CCR6^−^CD27^+^ phenotype. ScRNA-seq revealed their pro-inflammatory/B-Helper profile (*IL21, IFNG, TIGIT*, *CTLA4, NR3C1, CD109, KLRB1* and *CLEC2D)*. Autoreactive TCR clonotypes were restricted to the memory PD-1^+^CXCR5^−^ CD4 T cells. This subset was significantly increased in the blood of AIH patients and supported B cell differentiation through IL-21. Finally, we identified a specific phenotype (PD-1^+^CD38^+^CD27^+^CD127^−^CXCR5^−^) of CD4 T cells linked to disease activity and IgG response during AIH.

**Conclusions:** This work provides for the first time a deep characterization of rare circulating autoreactive CD4 T cells and the identification of their peripheral reservoir in AIH. We also propose a generic phenotype of pathogenic CD4 T cells related to AIH disease activity.

## Introduction

The main feature of autoimmune disorders is an abnormal reactivity of the adaptive immune system against self-antigens. The detection of the resulting self-reactive autoantibodies is one of the main criteria for the diagnosis of autoimmune diseases. Antibody production results from a crosstalk between cognate antigen-specific B cells and CD4 T cells in response to a specific antigen. The CD4 T follicular helper (TFH) cell population, characterized by the expression of the chemokine receptor CXCR5 and of the activation marker PD1, has been identified as the main subset of CD4 T cells responsible for B cell help in antibody responses^1–4^. However, the implication of TFH cells in human autoimmune activation is not yet well characterized, and whether those cells carry self-antigen-reactive TCR has not been studied in many human autoimmune diseases ^5–9^.

Autoimmune hepatitis (AIH) is a rare disease with an incidence range from 1 to 2 per 100 000 individuals in Europe. Like other autoimmune liver diseases (AILD), AIH is characterized by an immune attack targeting the liver and the accumulation of specific auto-antibodies (e.g. anti-SLA autoantibodies). A genetic predisposition linked to HLA class II alleles^10–12^ suggests a predominant role of CD4 T cell subsets in driving autoantibody accumulation and disease progression. Indeed, in AIH, CD4 T cells infiltrate the liver^10,13,14^, yet it is still debated whether those pathogenic CD4 T cells are related to TH17 cells^15–17^, TFH cells^7,8^ or TNF producing cells^14^. In fact, the molecular signature of self-antigen-specific CD4 T cells in AIH has remained elusive due to the difficulty to track such rare cells^10,18–21^, like in other AILD and many autoimmune disorders.

Recent studies have identified B-helper PD-1^+^CXCR5^−^ CD4 T cell populations, named T peripheral helper cell (TPH) or TH10 cells, which support antibody accumulation in rheumatoid arthritis (RA) through IL-21, or in systemic lupus erythematosus (SLE) through IL-10 production ^22–25^. Despite evidence for the accumulation of TPH or TH10 cells in their respective inflamed target tissue, it is still unknown whether they carry self-antigen-specific TCRs, a defining characteristic of autoreactive T cells. Christophersen et al. characterized gluten-specific CD4 T cells and proposed a generic distinct phenotype of CD4 T cell, based on the expression of PD-1 and the absence of CXCR5 expression, possibly involved in multiple autoimmune disorders such as systemic sclerosis, SLE and Celiac disease ^18^. However, although phenotypes are similar, the transcriptomic signatures of identified pathogenic PD-1^+^CXCR5^−^ CD4 T cells are different in RA^22^ and in SLE^23^, and not known in AIH.

In AIH, immunosuppressive treatments induce complete remission but more than 70% of patients relapse when treatment withdrawal is attempted, suggesting a persistence of pathogenic cells such as autoreactive CD4 T cells^10,26^. Moreover, the presence of anti-soluble liver antigen (SLA or SepSecs) autoantibodies in the blood of AIH patients is associated with significantly reduced overall survival^27^. This suggest a link between the adaptive immunity (autoreactive CD4 T cell response) and the prognosis of the AIH. Thus, there is an urgent clinical need to identify new therapeutic options targeting both the persistence and function of self-reactive T cells in AIH. Here, we have used *ex vivo* restimulation assays and 5’-end single-cell RNA-seq to define the surface phenotype, gene expression profile, and TCRαβ repertoire of a total of 546 self-reactive CD4 T cells in five distinct AIH patients with anti-SLA autoantibodies. We report that circulating autoantigen-specific CD4 T cells invariably carry a memory PD-1^+^CXCR5^−^ phenotype and a pro-inflammatory B-helper molecular and functional profile. Our study further identifies a specific T cell phenotypic signature which may be used as a blood biomarker of disease activity in all AIH patients.

## Methods

### Subject samples

All the patients eligible signed a written informed consent prior to inclusion and a bio-bank of samples from AIH patients (BioHAI) is maintained in Nantes University Hospital. All AIH patients had a simplified diagnostic score superior or equal to 6 according to the simplified scoring system for AIH of the international autoimmune hepatitis group (IAHG)^28^. All the patients received initially a standard immunosuppressive treatment protocol with corticosteroids (0.5-1 mg/kg). Azathioprine was added after two weeks (1-2 mg/kg) and monitored according to the tolerance of the patient. Corticosteroid treatment was then tapered until withdrawal. All the patients were screened for thiopurine S-methyltransferase (TPMT) genotyping before azathioprine administration. Untreated patients are new onset AIH patients enrolled at diagnosis prior any treatment initiation as previously described^13^. Treated / Active AIH patients are under standard treatment but do not normalize the transaminases (AST and ALT) or the serum IgG levels or present an active interface hepatitis in the liver biopsy. Treated / Complete remission AIH patients are defined biochemically by a normalization of the transaminases and the IgG levels, according to the most recent European clinical practice guidelines on AIH^29^.

### Peptide re-stimulation assay

Peripheral blood mononuclear cells (PBMCs) from six SLA-pos (seropositive) and six SLA-neg (seronegative) AIH patients were tested for their reactivity against SLA and MP65 peptides (Candidas Albicans / C.Alb antigen used as positive control: common antigen with a TH17 signature). For the CD154 (CD40L) expression assay, 10 to 20×10^6^ PBMCs (at a final concentration of 10×10^6^/mL) were stimulated for 3h at 37°C with 5µg/mL of synthesized peptide pools (20 amino acids in length with a 12 amino acid overlap; Synpeptide, China) spanning all of the Soluble Liver Antigen (SepSecs) sequences [SLA (p1-p53)] or 0.6 nmol/mL PepTivator^R^ Candida albicans MP65 (peptides pools of 15 amino acids length with 11 amino acid overlap, Miltenyi Biotec) in 5% human serum RPMI medium in the presence of 1µg/ml anti-CD40 (HB14, Miltenyi Biotec). After 3 hrs of specific peptide stimulation, PBMCs were first labeled with PE-conjugated anti-CD154 (5C8, Miltenyi Biotec) and CD154^+^ cells were then enriched using anti-PE magnetic beads (Miltenyi Biotec). A 1/10th fraction of non-enriched cells was saved for frequency determination. Frequency was calculated with the formula F = n/N, where n is the number of CD154 positive cells in the bound fraction after enrichment and N is the total number of CD4+ T cells (calculated as 10 × the number of CD4+ T cells in 1/10th non-enriched fraction that was saved for analysis). After enrichment, cells were stained with PerCP-Cy5.5 anti-CD4 (RPA-T4, Biolegend), Alexa Fluor 700 anti-CD3 (SK7, Biolegend), APC-Cy7 anti-CD45RA (HI100, Biolegend), Alexa Fluor 647 anti-CD185 (CXCR5, J252D4, Biolegend), Alexa Fluor 488 anti-CD196 (CCR6, G034E3, Biolegend), PE-Cy7 anti-PD-1 (EH12.2H7, Biolegend) antibodies.

### Flow cytometry and cell sorting

PerCP-Cy5.5, PE and Brilliant violet 711 anti-CD4 (RPA-T4, Biolegend), Alexa Fluor 700 and PerCP-Cy5.5 anti-CD3 (SK7, Biolegend), APC-Cy7 and Brilliant violet 605 anti-CD45RA (HI100, Biolegend), Alexa Fluor 647 anti-CD185 (CXCR5, J252D4, Biolegend), Alexa Fluor 488 anti-CD196 (CCR6, G034E3, Biolegend), PE-Cy7 anti-PD-1 (EH12.2H7, Biolegend), PerCP-Cy5.5 anti-CD38 (HB-7, Biolegend), Brilliant Violet 421 anti-CD27 (M-T271, Biolegend), Brilliant Violet 605 anti-TIGIT (A15153G, Biolegend), PerCP-Cy5.5 anti-CD49d (9F10, Biolegend), PE anti-CD127 (A019D5, Biolegend), APC-Cy7 anti-CD8 (SK1, Biolegend), Brilliant Violet anti-CD278 (ICOS, DX29, BD Biosciences), Alexa Fluor 488 and Brilliant violet 605 anti-CD19 (HIB19, Biolegend) antibodies were used for surface staining after LIVE/DEAD™ Fixable Aqua Dead Cell staining (ThermoFisher scientific). Briefly, PBMCs were incubated 20 minutes with a mix of antibodies and then washed prior analysis or cell sorting on BD FACSLSRII or BD FACSAriaII.

PE anti-CD152 (CTLA4, BNI3, BD Biosciences), Alexa Fluor 700 anti-IFNG (B27, BD Biosciences) and PE anti-IL21 (3A3-N2.1, BD Biosciences) antibodies were used for intra-cellular staining by using the Fixation/Permeabilization Solution Kit (BD Cytofix/Cytoperm™, BD Biosciences). For *in vitro* intracellular cytokine staining, cells were restimulated with 100 ng/mL phorbol 12-myristate 13-acetate and 1 µg/mL ionomycin in the presence of 10 mg/mL Brefeldin-A for 4 hours at 37ºC prior surface and intra-cellular staining.

### Single cell RNA sequencing and analysis (scRNAseq)

First, CD154^+^ memory CD4 T cells were sorted on BD FACSAriaII, one cell per well, in 96-well plates containing specific lysis buffer at the CRTI, Nantes. Plates were immediately frozen for storage at −80°C, and sent on dry ice to the Genomics core facility of CIML, Marseille, for further generating scRNAseq libraries with the FB5P-seq protocol as described^30^. Briefly, mRNA reverse transcription (RT), cDNA 5’-end barcoding and PCR amplification were performed with a template switching (TS) approach. After amplification, barcoded full-length cDNA from each well were pooled for purification and library preparation. For each plate, an Illumina sequencing library targeting the 5’-end of barcoded cDNA was prepared by a modified transposase-based method incorporating a plate-associated i7 barcode. Resulting libraries had a broad size distribution, resulting in gene template reads covering the 5’-end of transcripts from the 3^rd^ to the 60^th^ percentile of gene body length on average. As a consequence, sequencing reads cover the whole variable and a significant portion of the constant region of the TCRα and TCRβ expressed mRNAs, enabling assembly and reconstitution of TCR repertoire from scRNAseq data.

Cells were analyzed in three distinct scRNAseq experiments (**Extended Table 1**). Libraries prepared with the FB5P-seq protocol were sequenced on Illumina NextSeq550 platform with High Output 75 cycles flow cells, targeting 5×10^5^ reads per cell in paired-end single-index mode with the following configuration: Read1 (gene template) 67 cycles, Read i7 (plate barcode) 8 cycles, Read2 (cell barcode and Unique Molecular Identifier) 16 cycles.

We used a custom bioinformatics pipeline to process fastq files and generate single-cell gene expression matrices and TCR sequence files, as described^30^. Briefly, the pipeline to obtain gene expression matrices was adapted from the Drop-seq pipeline^31^, relied on extracting the cell barcode and UMI from Read2 and aligning Read1 on the reference genome using STAR and HTSeqCount. For TCR sequence reconstruction, we used Trinity for *de novo* transcriptome assembly for each cell based on Read1 sequences, then filtered the resulting isoforms for productive TCR sequences using MigMap, Blastn and Kallisto. Briefly, MigMap was used to assess whether reconstructed contigs corresponded to a productive V(D)J rearrangement and to identify germline V, D and J genes and CDR3 sequence for each contig. For each cell, reconstructed contigs corresponding to the same V(D)J rearrangement were merged, keeping the largest sequence for further analysis. We used Blastn to align the reconstructed TCR contigs against reference sequences of constant region genes, and discarded contigs with no constant region identified in-frame with the V(D)J rearrangement. Finally, we used the pseudoaligner Kallisto to map each cell’s FB5P-seq Read1 sequences on its reconstructed contigs and quantify contig expression. In cases where several contigs corresponding to the same TCR chain had passed the above filters, we retained the contig with the highest expression level.

Quality control was performed on each scRNAseq batch independently to remove poor quality cells. Cells with less than 250 genes detected were removed. We further excluded cells with values below 3 median absolute deviations (MADs) from the median for UMI counts, for the number of genes detected, or for ERCC accuracy, and cells with values above 3 MADs from the median for ERCC transcript percentage, as described^30^. The number of cells passing quality control, mean sequencing depth, and mean gene detection for each batch is shown in **Extended Table 1**.

All cells passing quality control were analyzed together with custom scripts in the R programming language (available upon request). For each cell, gene expression UMI count values were log normalized with Seurat v3.0.0.9^32^ *NormalizeData* with a scale factor of 10,000. Four thousand variable genes were identified with Seurat *FindVariableFeatures*, of which TCR coding genes were removed, yielding 3919 variable genes for further analyses. After data centering with Seurat *ScaleData*, principal component analysis was performed on 3919 variable genes with Seurat *RunPCA*, and embedded in two-dimensional tSNE with Seurat *RunTSNE* on 40 principal components. Differentially expressed genes between C.Alb-specific and SLA-specific T cells were identified with Seurat *FindAllMarkers* which computes a likelihood-ratio test for single-cell gene expression (test.use=’bimod), with an adjusted p-value cutoff of 0.05. Gene expression heatmaps and plots showing PCA or tSNE embeddings colored by gene expression levels or metadata were plotted with custom functions and ggplot2 *ggplot*.

Following *in silico* reconstruction of TCR sequences from scRNAseq reads, functional variable TCRα or TCRβ sequences were recovered for 79%, 75% and 54% of cells passing gene expression quality control in the three scRNAseq experiments, respectively (**Extended Table 1**). Nucleotide sequences were submitted to IMGT HighV-QUEST analysis^33^ to identify V and J genes, CDR3 junction sequence, and other TCR sequence features. We further processed the IMGT HighV-QUEST output summary table in Microsoft Excel. To identify recurrent clonotypes, we first defined TCRα and TCRβ clonotypes independently, based on identical CDR3 juntion amino acid sequences. Cells with both TCRα and TCRβ sequences were grouped into TCRαβ clonotypes if they were of the same TCRα and TCRβ clonotypes. Cells for which only one of the two sequences was available were joined to a TCRαβ clonotype based only on its TCRα or TCRβ clonotype identity. Clonotype identity for each cell was added as a metadata to the Seurat scRNAseq dataset for visualization in tSNE embeddings.

### TCR sequencing

100×10^3^ PD-1^+^CD45RA^−^CXCR5^−^CCR6^−^, PD-1^−^CD45RA^−^CXCR5^−^CCR6^+^ and PD-1^−^CD45RA^−^CXCR5^−^ CCR6^−^ CD4 T cell subsets were sorted on a BD FACSAriaII. gDNA were extracted with NucleoSpin® Blood kit (Macherey-Nagel) and TCR sequencing was performed by Adaptive Biotechnologies^R^ (Seattle, USA).

### B cell co-culture assay

Total B cells (CD19^+^CD4^−^CD3^−^), PD-1^+^CXCR5^−^CD45RA^−^ and PD-1^−^CXCR5^−^CD45RA^−^ CD4 T cells were sorted on BD FACSAriaII. Sorted cell subsets were co-cultured at a ratio 1:1 in 200µL RPMI/10% FBS and stimulated with CytoStim™ (Miltenyi Biotec). After 7 days’ culture, supernatants were collected and total IgG measured by ELISA (eBioscience). Plasmablasts (CD19^+^CD38^+^CD27^+^) percentage was analyzed by flow cytometry. For blocking experiments, 20µg/mL IL-21 R Fc Chimera (R&D Systems) was used.

### Unsupervised flow cytometry analysis

Flow cytometry data from 22 patients (11 active AIH and 11 NASH) were generated the same day and first analyzed using FlowJo 10.6.0 (TreeStar Inc) to extract the same number of viable lymphoid CD3+ cells (27.10^3^ per patients; total of 594.10^3^). New FCS folders were generated with identical CD3+ T cell number to performed unsupervised clustering analysis. FlowSOM package on R was used to generate 400 clusters. Metaclustering was performed by using WGCNA package. Heat-maps were generated by using complex heat map package. Wilcoxon test was used to identify significant variation between active AIH and NASH patients.

### Statistical analysis

Statistical comparisons were performed as indicated in figure legends. P value <0.05 after adjustment were considered significant.

### Accession codes

The single-cell RNA-seq data generated in the current study are available in the Gene Expression Omnibus database under accession code GSExxxxxx.

## Results

### SLA-specific CD4 T cells are detectable in the blood of seropositive (anti-SLA+) AIH patients and present a memory PD-1^+^CXCR5^−^ phenotype

To directly identify autoreactive CD4 T cells based on their antigen specificity, we performed an *ex vivo* short stimulation assay on PBMC with a pool of antigen-derived overlapping peptides, followed by an enrichment step of antigen-specific CD154^+^ T cells^34–36^ (**Extended Data Figure 1)**. We analyzed AIH patients with (SLA-pos) and without (SLA-neg) anti-SLA autoantibodies and detected SLA-specific CD4 T cells only in SLA-pos patients, whereas control C.Alb-specific CD4 T cell counts were similar in both groups (Figure 1a and b and Table 1). The SLA-specific CD4 T cell population frequency was stable in longitudinal analysis of two SLA-pos AIH patients (**Extended Data Figure 1**).

**Table 1.** SLA-neg and SLA-pos patients’ characteristic

| **Patients** | **#006** | **#027** | **#030** | **#073** | **#085** | **#106** | **#004** | **#018** | **#028** | **#051** | **#128** | **#140** |
| --- | --- | --- | --- | --- | --- | --- | --- | --- | --- | --- | --- | --- |
| **Age** | 54 | 55 | 68 | 66 | 72 | 38 | 33 | 32 | 54 | 30 | 27 | 20 |
| **Sex** | F | F | M | F | F | F | F | F | M | F | F | F |
| **AIH score** | 7 | 7 | 6 | 7 | 8 | 7 | 7 | 6 | 6 | 7 | 7 | 6 |
| **Disease** |  |  |  |  |  |  |  |  |  |  |  |  |
| **duration** | 12 | 4 | 4 | 10 | 2 | 0 | 15 | 11 | 10 | 20 | 10 | 5 |
| **(years)** |  |  |  |  |  |  |  |  |  |  |  |  |
| **status at** | complete | Active under | complete | complete | complete | untreated / | complete | Active under | Active under | complete | complete | complete |
| **experiment** | remission | treatment* | remission | remission | remission | diagnostic | remission | treatment* | treatment* | remission | remission | remission |
| **time point** |  |  |  |  |  |  |  |  |  |  |  |  |
| **anti-SLA** | SLA-neg | SLA-neg | SLA-neg | SLA-neg | SLA-neg | SLA-neg | SLA-pos | SLA-pos | SLA-pos | SLA-pos | SLA-pos | SLA-pos |
| **Treatment** | neoral | imurel | cellcept | imurel | imurel | none | imurel | tacrolimus | imurel | imurel | imurel | UCDA/imurel |

\* AST or ALT or IgG>ULN or active interface hepatitis on biopsy

**Figure 1:**
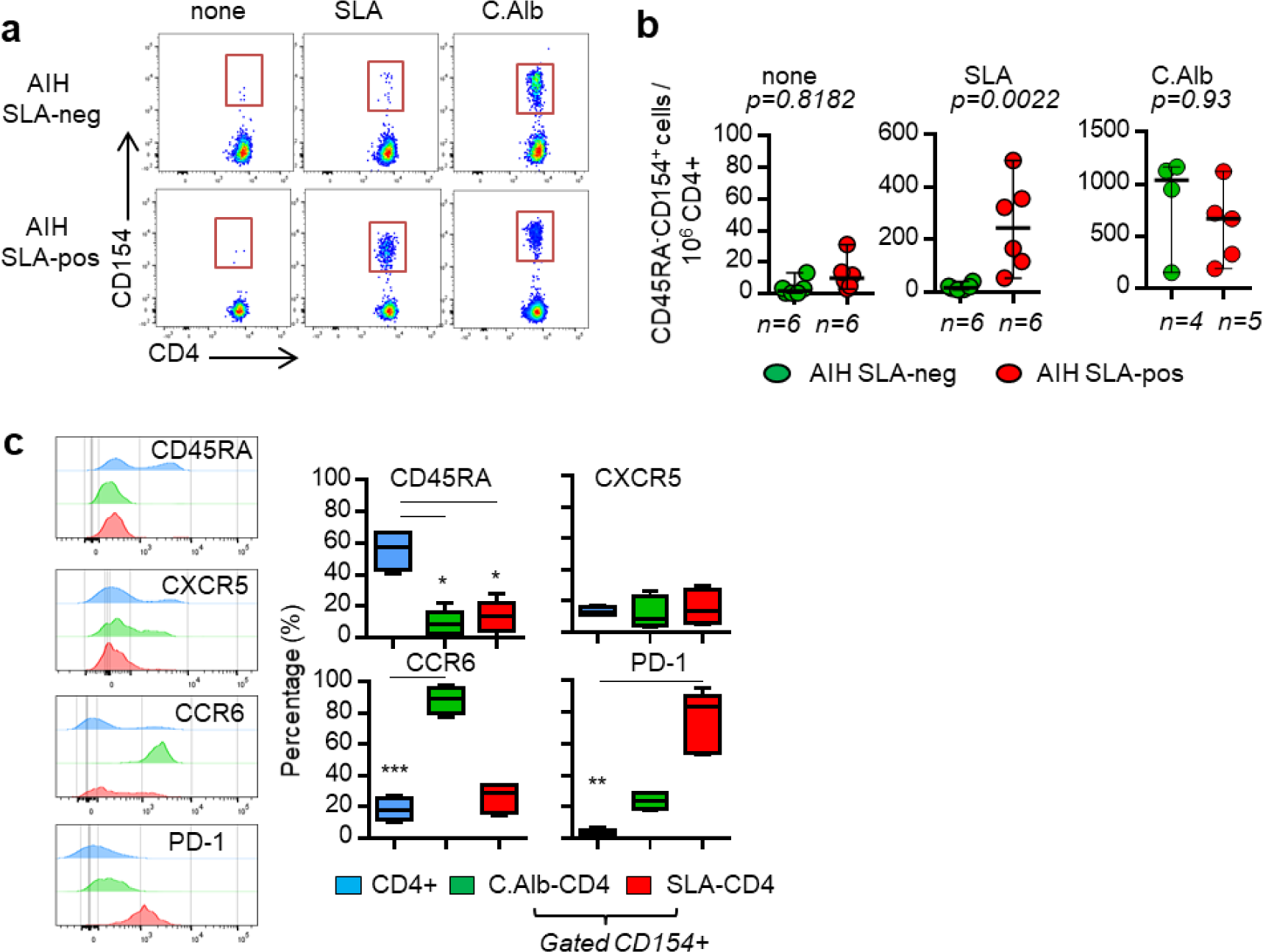
Memory PD-1^+^CXCR5^−^ SLA-specific CD4 T cells in the blood of seropositive (anti-SLA+) AIH patients. **a**, Dot plot of the CD154 expression on CD4 T cells, after short stimulation with overlapping peptides, derived from the antigen SLA or MP65 (Candida Albicans; C.Alb), or in control condition (none), in a seronegative (anti-SLA-neg) and a seropositive (anti-SLA-pos) AIH patient. **b**, Frequency of CD154^+^CD4^+^CD45RA^−^ cells per million total CD4 T cells from 7 seronegative and 6 seropositive AIH patients. **c**, Histogram and frequency of CD45RA, CXCR5, CCR6 and PD-1 expression on total CD4 T cells (blue), CD154^+^CD4+ T cells after stimulation with MP65 (C.Alb-CD4; green) or SLA peptides (SLA-CD4; red). Comparisons were performed using the Mann-Whitney U test (**b**) or the Kruskal-Wallis test and Dunn’s Multiple Comparison Test (**c**). *: p<0,05; **: p<0,01; ***: p<0,001.

Both SLA- and C.Alb-specific CD4 T cells exhibited a memory phenotype but differed for the expression of CCR6 and PD-1 (Figure 1c and **Extended Data Figure 1**). C.Alb-specific CD4 T cells were PD-1^−^ CXCR5^−^CCR6^+^, consistent with their expected TH17 phenotype^36–38^. Autoreactive SLA-specific CD4 T cells were PD-1^+^CXCR5^−^CCR6^−^ and express CD27 (**Extended Data Figure 1**). Neither SLA- nor C.Alb-specific CD4 T cells showed high CXCR5 expression, suggesting they were distinct from circulating TFH CD4 T cells, classically involved in the B cell differentiation process^1,39^.

### scRNA-seq analysis reveals a TPH-like profile of auto-reactive CD4 T cells

Next we submitted *ex vivo* activated C.Alb-specific and SLA-specific memory CD4 T cells from 5 SLA-pos AIH patients to FACS-based 5’-end single-cell RNA-seq analysis for parallel analysis of gene expression and TCRαβ sequence^30^ (Figure 2a). Unsupervised analysis of the single-cell gene expression profiles revealed a clear distinction between the two antigen-specific memory T cell subsets (Figure 2b). A total of 211 genes were differentially expressed between C.Alb-specific and SLA-specific CD4 T cells (Figure 2c and d and **Extended Table 2**). The molecular profiles of C.Alb-specific and SLA-specific CD4 T cells were robust and stable in a longitudinal analysis of samples from one patient collected at 3 time points over a 6-month period (Figure 2c).

**Figure 2:**
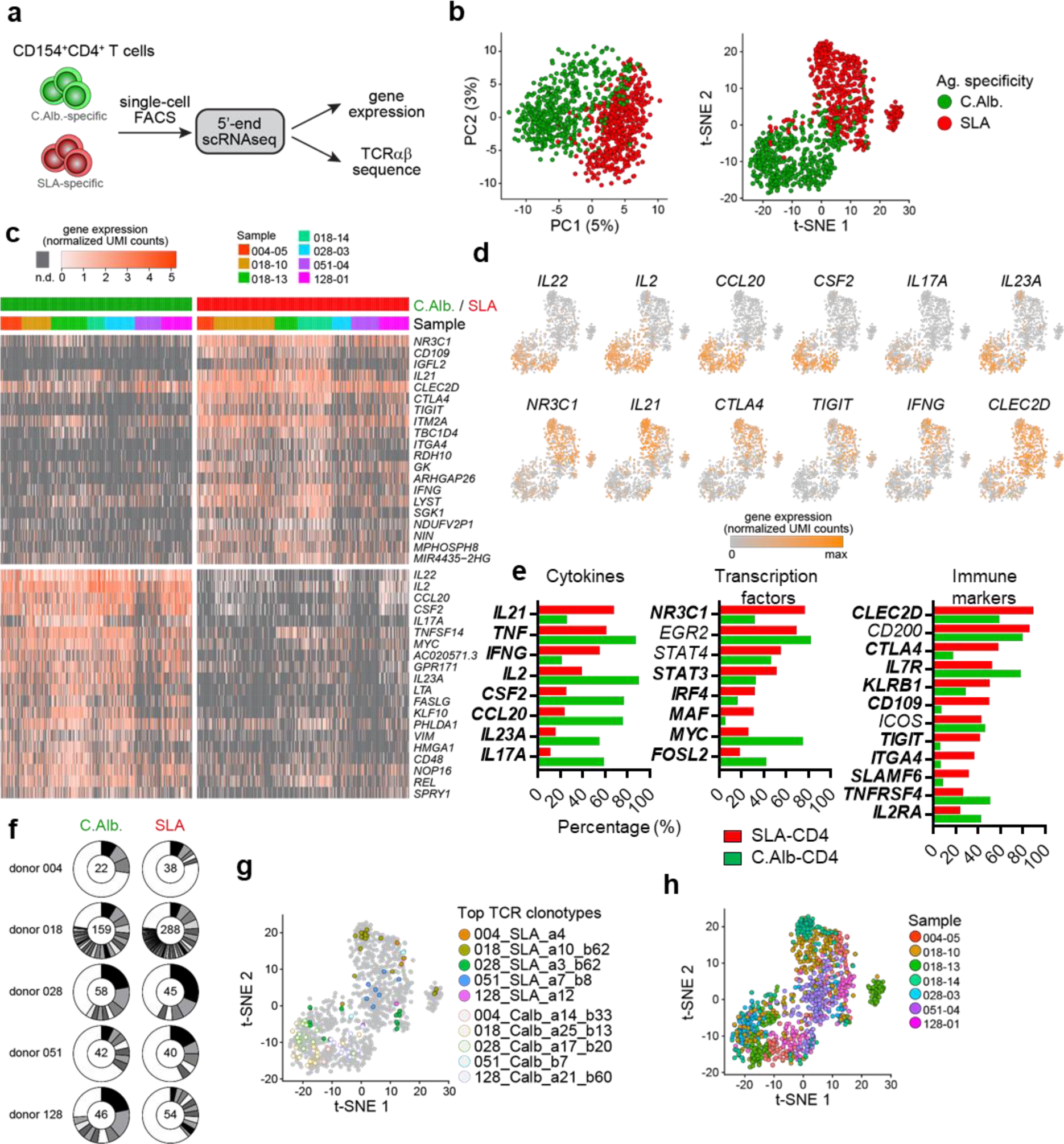
scRNAseq analysis reveal a TPH-like profile of auto-reactive CD4 T cells. **a**, Experimental design for scRNAseq analysis of antigen-specific T cells. **b**, PCA and Unsupervised dimensionality reduction (tSNE) of single-cell transcriptomes of n=546 SLA-specific (red) and n=493 C.Alb.-specific (green) CD4 T cells isolated from peripheral blood of 5 distinct AIH patients. **c**, heat map representation of the top 20 genes differentially expressed in SLA-specific (red) and C.Alb.-specific (green) CD4 T cells. **d**, Single-cell gene expression tSNE plots showing expression levels of C.Alb.-specific marker genes (top row) and SLA-specific marker genes (bottom rows), as indicated. **e**, Frequency of SLA-specific (red) and C.Alb.-specific (green) CD4 T cells expressing genes related to cytokines, transcription factors or immune markers. Bold genes are significantly different between groups. **f**, Distribution of TCRαβ clones among each patient’s antigen-specific T cells. Numbers indicate the number of single cells analyzed per patient. Black and grey sectors indicate the proportion of TCRαβ clones (clonotype expressed by ≥ 2 cells) within single-cells analyzed (white sector: unique clonotypes). **g**, Same tSNE plot presented in Figure 2b with the Top TCRαβ clones identified (colored dots) per patients overlaid on the total dataset (full circle, SLA-specific; empty circle, C.Alb.-specific). **h**, Same tSNE plot presented in Figure 2b with each patients represented.

One advantage of the short restimulation assay is to allow transcriptomic characterization of the cytokine expression profile of antigen-specific CD4 T cells^34^ (Figure 2d and e). C.Alb-specific CD4 T cells showed a typical TH17 cytokine profile^37,40^ with high expression of *IL17A*, *IL23A*, *CCL20*, *CSF2* (GM-CSF)*, IL2* and *TNF* among other genes. C.Alb-specific CD4 T cells were characterized by the expression of the transcription factors *FOSL2, MYC* and *EGR2*, characteristic genes expressed in the TH17 lineage^37,40^ (Figure 2d and e and **Extended Table 2**). C.Alb-specific CD4 T cells also expressed genes encoding the surface immune markers *CD200, ICOS, IL7R* (CD127) and *TNFRSF4* (OX40). By contrast, restimulated circulating autoreactive SLA-specific CD4 T cells had a distinct B-helper (*IL21*) and pro-inflammatory (*IFNG and TNF*) gene signature (Figure 2d and e and **Extended Table 2)**. This B-helper profile was also conferred by the expression of the transcription factor *MAF* and by the expression of genes encoding surface immune markers such as *ICOS, TIGIT, CTLA4* and *SLAMF6*, which were previously found to be elevated on both TFH and TPH cells^22,41^. SLA-specific CD4 T cells did not express CXCR5 (Figure 1) and expressed *IFNG* suggesting their molecular signature was closer to the reported TPH signature (*CD200*, *IL21, IFNG, ICOS, TIGIT, CTLA4, ITM2A, SLAMF6* and *MAF*) than to a TFH signature^22,24,25^. However, SLA-specific CD4 T cells specifically expressed some additional genes not previously reported in studies of human autoimmune TPH or TFH cells: the transcription factor *NR3C1*, a glucocorticoid receptor involved in immune-regulation processes^42^; *CD109*, an inhibitor of the TGF-β pathway^43,44^ and also involved in the regulation of inflammation^45^; *CLEC2D* and *KLRB1*, involved in the regulation of the innate immune response^46^; and *ITGA4* (CD49d) and *ITGB1*, which together form the complex VLA-4. Thus, autoreactive CD4 T cells in AIH share a transcriptomic signature with the previously reported TPH cell population but differ for the expression of additional immune regulatory genes.

With the 5’-end scRNA-seq method we were able to reconstruct paired variable TCRα/TCRβ sequences *in silico* and link them to the transcriptome of antigen-specific CD4 T cells (Figure 2a). In all patients, SLA- and C.Alb-specific CD4 T cells were polyclonal, yet we consistently identified some highly frequent TCRαβ clonotypes indicative of antigen-specific clonal expansions (Figure 2f). We found no evidence for public TCR rearrangements in SLA- and C.Alb-specific CD4 T cells from these five AIH patients (**Extended Table 3**). T cells from the same TCRαβ clonotype were transcriptionally diverse within the overall molecular profile associated with their antigen-specificity (Figure 2g), which was conserved across patients despite some detectable sample-related batch effects (Figure 2h).

### The PD-1^+^CXCR5^−^ CD4 T cell population is the reservoir of auto-reactive CD4 T cells in AIH

Given the B-helper molecular profile and PD-1^+^ CXCR5^−^ surface phenotype of *ex vivo* activated SLA-specific CD4 T cells, we set out to more precisely identify the phenotypic niche of autoreactive CD4 T cells in the peripheral blood of AIH patients. Bulk TCRβ sequencing was performed on sorted memory PD-1^+^CXCR5^−^CCR6^−^, PD-1^−^CXCR5^−^CCR6^+^ and PD-1^−^CXCR5^−^CCR6^−^ CD4 T cell subsets sampled at three different time points from the same patient (**Extended Data Figure 2)**. SLA-specific TCRβ clonotypes identified at the single-cell level were then tracked in the bulk datasets and found to be restricted to the PD-1^+^CXCR5^−^CCR6^−^ CD4 T cell population (Figure 3a). In contrast, C.Alb-specific TCRβ clonotypes were restricted to the PD-1^−^CXCR5^−^CCR6^+^ CD4 T cell population (Figure 3b). The relative frequency of TCRβ clonotypes was stable in longitudinal sampling across 6 to 8 months, both for SLA- and C.Alb-specific CD4 T cells (**Extended Data Figure 2b and c**). These results clearly demonstrate that the PD-1^+^CXCR5^−^ CD4 T cell population is the major peripheral reservoir of autoreactive CD4 T cells in AIH.

**Figure 3.**
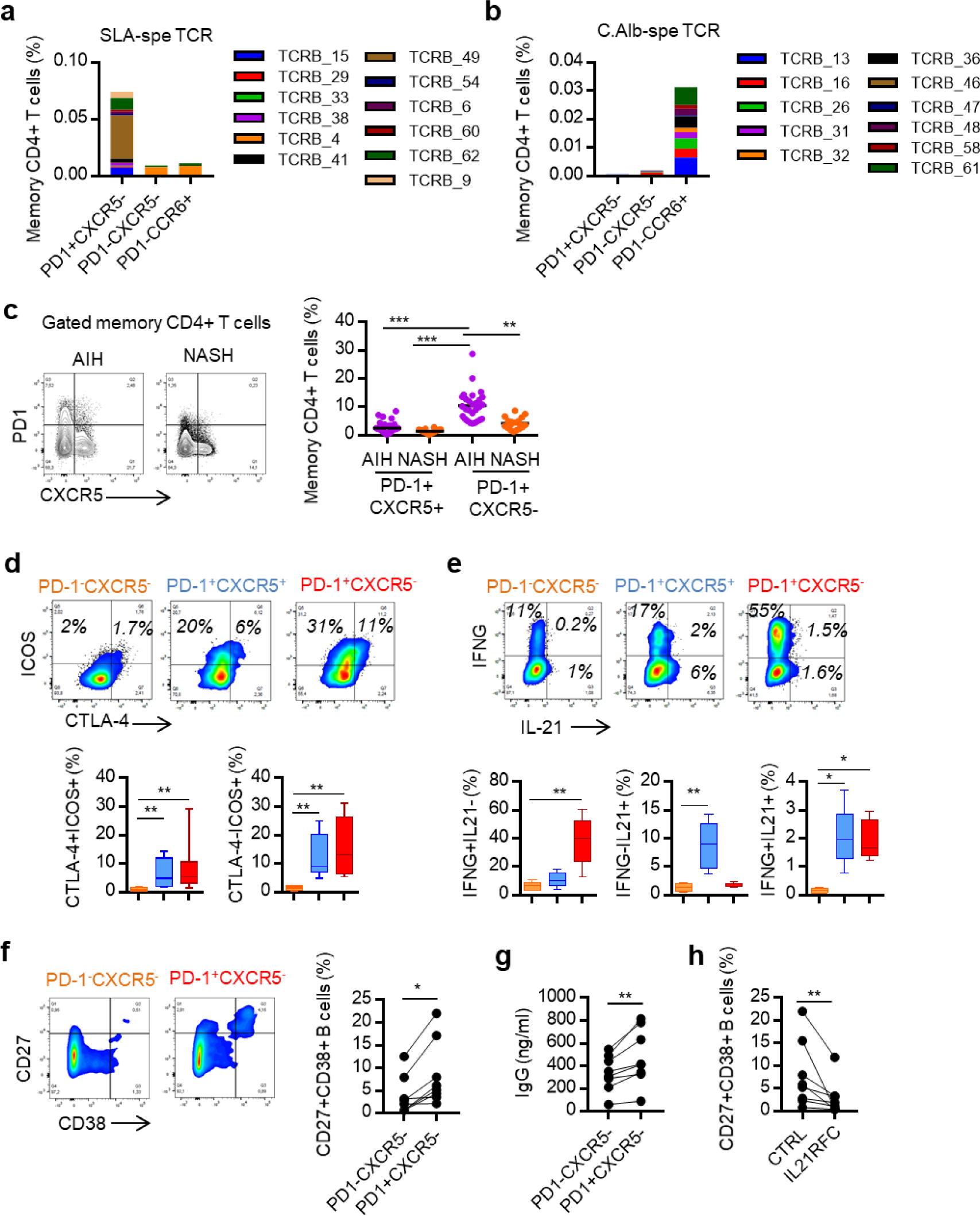
The PD-1^+^CXCR5^−^ CD4 T cell population is the reservoir of auto-reactive CD4 T cells with a TPH-like phenotype. **a** and **b**, Frequency of the major TCRβ clonotypes of SLA-CD4 T cells (**a**) and the major TCRβ clonotypes of C.Alb-CD4 T cells (**b**) identified in the donor 018 (Figure 2) within PD-1^+^CXCR5^−^(CCR6^−^), PD-1^−^CCR6^+^(CXCR5^−^) and PD-1^−^CXCR5^−^(CCR6^−^) memory CD4 T cells isolated from the same patient at three different time points. Data presented are the three time points aggregated. **c**, Dot plot and graph of the CD45RA^−^PD-1^+^CXCR5^−^ and the CD45RA^−^PD-1^+^CXCR5^+^ CD4+ T cell populations in active AIH (n=28) and NASH (n=18) patients. **d** and **e**, Percentage of ICOS and CTLA4 expression (n=8) (**d**) and of IFNG and IL21 expression (n=5) (**e**) in PD-1^+^CXCR5^−^ (red), PD-1^+^CXCR5^+^ (blue) and PD-1^−^CXCR5^−^ (orange) memory CD4 T cell subsets in AIH patients. **f** and **h**, Percentage of plasma cells (**f**; CD27^+^CD38^+^) after a 7 day co-culture assay with autologous B cells and memory CD4 T cell subsets with IL21RFC or without (CTRL) (**h**). **g**, Concentration of IgG after a 7 day co-culture assay with autologous B cells and memory CD4 T cell subsets. **f-h**, each point represents a duplicate experiment per patients (patients, n=8). Comparisons were performed using the Kruskal-Wallis test and Dunn’s Multiple Comparison Test (**c, d** and **e**) or the Wilcoxon matched-pairs signed rank test (**f**, **g** and **h**). *: p<0,05; **: p<0,01; ***: p<0,001; ****: p<0,0001.

### Memory PD-1^+^CXCR5^−^ CD4 T cells are enriched in the blood of AIH patients

We compared the immune phenotype of PBMC between patients with an active AIH (n=28, treated or not) and patients with Non-Alcoholic SteatoHepatitis (NASH) with no sign of autoimmunity (n=18) (Figure 3c and Table 2). The proportion of PD-1^+^CXCR5^−^ cells amongst the memory CD4 T cell population was increased in AIH patients and exceeded that of the PD-1^+^CXCR5^+^ TFH cell population (Figure 3c).

**Table 2.**
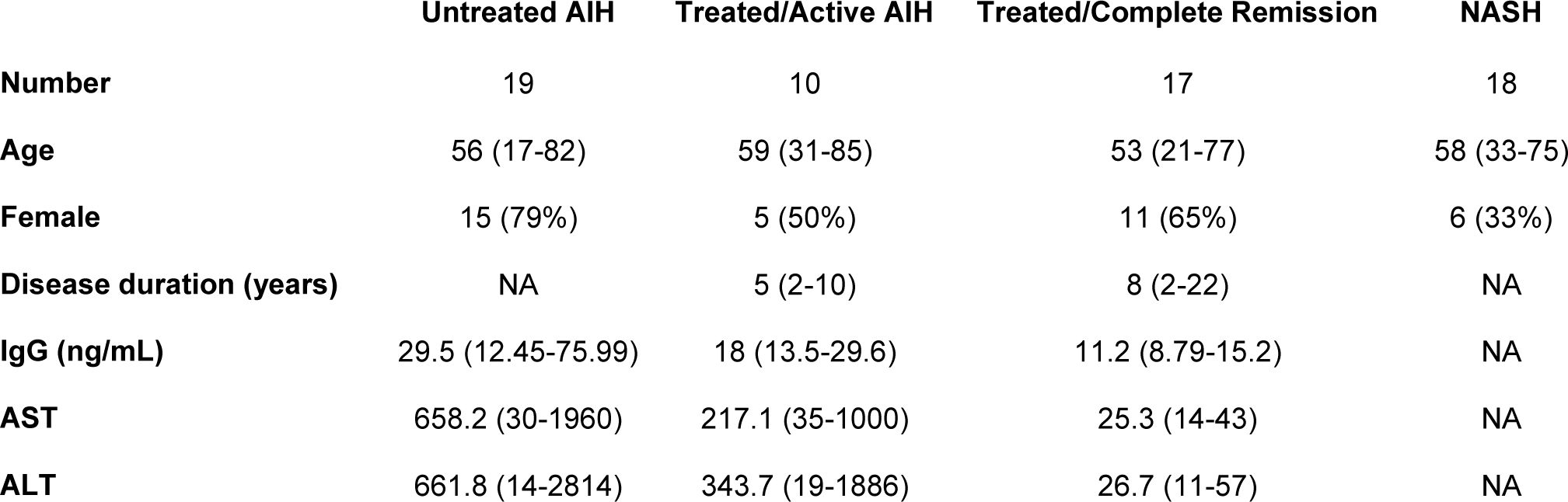
Clinical characteristics of AIH and NASH patients

|  | Untreated AIH | Treated/Active AIH | Treated/Complete Remission | NASH |
| --- | --- | --- | --- | --- |
| **Number** | 19 | 10 | 17 | 18 |
| **Age** | 56 (17-82) | 59 (31-85) | 53 (21-77) | 58 (33-75) |
| **Female** | 15 (79%) | 5 (50%) | 11 (65%) | 6 (33%) |
| **Disease duration (years)** | NA | 5 (2-10) | 8 (2-22) | NA |
| **IgG (ng/mL)** | 29.5 (12.45-75.99) | 18 (13.5-29.6) | 11.2 (8.79-15.2) | NA |
| **AST** | 658.2 (30-1960) | 217.1 (35-1000) | 25.3 (14-43) | NA |
| **ALT** | 661.8 (14-2814) | 343.7 (19-1886) | 26.7 (11-57) | NA |

### The PD-1^+^CXCR5^−^ CD4 T cell population has a TPH-like phenotype and function in AIH

Next, we established the phenotypic link between SLA-specific CD4 T cells and the peripheral PD-1^+^CXCR5^−^ memory CD4 T cell population in a group of AIH patients independent of their anti-SLA antibody serology. Recent studies have reported phenotypic and functional similarities between blood memory PD-1^+^CXCR5^−^ CD4 TPH cells and PD-1^+^CXCR5^+^ CD4 TFH cells^22^. In AIH patients, both PD-1^+^CXCR5^−^ and PD-1^+^CXCR5^+^ memory CD4 T cells were TIGIT^high^CD127^low^CD49d^high^ and expressed ICOS and CTLA4, similar to SLA-specific CD4 T cells (Figure 3d and **Extended Data Figure 3**). Yet we found major differences regarding their capacity to produce IFNγ and IL-21 upon *ex vivo* restimulation. In PD-1^+^CXCR5^−^ CD4 T cells, 38.3 +/− 17.3 % cells produced IFNγ, and more than half of IL-21 producing cells also produced IFNγ. In contrast, only 10.8 +/− 5.2 % of PD-1^+^CXCR5^+^ TFH produced IFNγ and most IL-21 producing TFH cells did not produce IFNγ (Figure 3e). These results are consistent with the pro-inflammatory profile of pathogenic PD1^+^CXCR5^−^ CD4 T cells and SLA-specific CD4 T cells. To test the B helper capacity of PD-1^+^CXCR5^−^ CD4 T cells from AIH patients (n=8; one experiment per patient), we sorted PD-1^+^CXCR5^−^ and PD-1^−^CXCR5^−^ memory CD4 T cells and co-cultured them with autologous B cells in the presence of a super-antigen (Figure 3f-h). Memory PD-1^+^CXCR5^−^ CD4 T cells from AIH patients were more potent than PD-1^−^CXCR5^−^ CD4 T cells at inducing the differentiation of B cells into CD27^+^CD38^+^ plasmablasts (Figure 3f) and IgG secretion (Figure 3g). Blocking soluble IL-21 with decoy IL-21R in those co-cultures drastically reduced the proportion of differentiated plasmablasts (Figure 3h), suggesting that the B-helper capacity of PD1^+^ CXCR5^−^ CD4 T cells in AIH patients was mostly IL-21-dependent.

### PD-1 and CD38 as potential blood markers of an active AIH

In a parallel effort to determine the clinical relevance of tracking the autoreactive CD4 T cell population and to more precisely delineate its specific phenotype, we performed 11-color flow cytometry on peripheral blood T cells from 11 patients with an active AIH versus 11 patients with Non-Alcoholic SteatoHepatitis (NASH) (Table 2), monitoring markers identified in the scRNA-seq data or in previous phenotypic observations (Figure 1, 2 and 3), with a focus on activation and differentiation markers (CD38, CD127 and CD27). Unsupervised analysis of these data with self-organizing maps **(Extended Data Figure 4**) and hierarchical clustering (**Extended Data Figure 5**) identified 14 phenotypic T cell metaclusters (Figure 4a). Three metaclusters were significantly increased in AIH patients compared to NASH patients (Figure 4b). Those three clusters were respectively memory CD4^+^, CD8^+^ and double negative T cells, and all shared the PD-1^+^CD38^+^CXCR5^−^CCR6^−^CD127^−^CD27^+^ phenotype (Figure 4c and **Extended Data Figure 6**). With a focus on the CD4 T cell population, the frequency of PD-1^+^CD38^+^CXCR5^−^CCR6^−^CD127^−^CD27^+^ memory CD4 T cells was significantly high in AIH patients with active disease compared not only to NASH patients, but also to AIH patients in complete remission under treatment (Figure 4d and **Extended Data Figure 7**). This latter difference was not observed for the frequency of the broader PD1^+^CXCR5^−^ memory CD4 T cell subset (Figure 4e). Moreover, with ROC curve analyses, we observed that only the frequency of PD1^+^CD38^+^CXCR5^−^CCR6^−^CD127^−^CD27^+^ memory CD4 T cells could discriminate active AIH from AIH patients in complete remission under treatment (AUC=0.73; p=0.08); this was not possible with the frequency of global PD1^+^CXCR5^−^ memory CD4 T cells (AUC=0.62; p=0.19) (Figure 4f and g). Finally, and Interestingly, the frequency of PD1^+^CD38^+^CXCR5^−^CCR6^−^CD127^−^CD27^+^ memory CD4 T cells was better correlated to the serum IgG level than the frequency of the PD1^+^CXCR5^−^ memory CD4 T cell subset (Figure 4h and i). These analyses suggested that tracking peripheral blood CD4 T cells expressing this combination of markers could be used as a specific biomarker of AIH activity, and should be evaluated for clinical use in AIH and in other autoimmune disorders.

**Figure 4.**
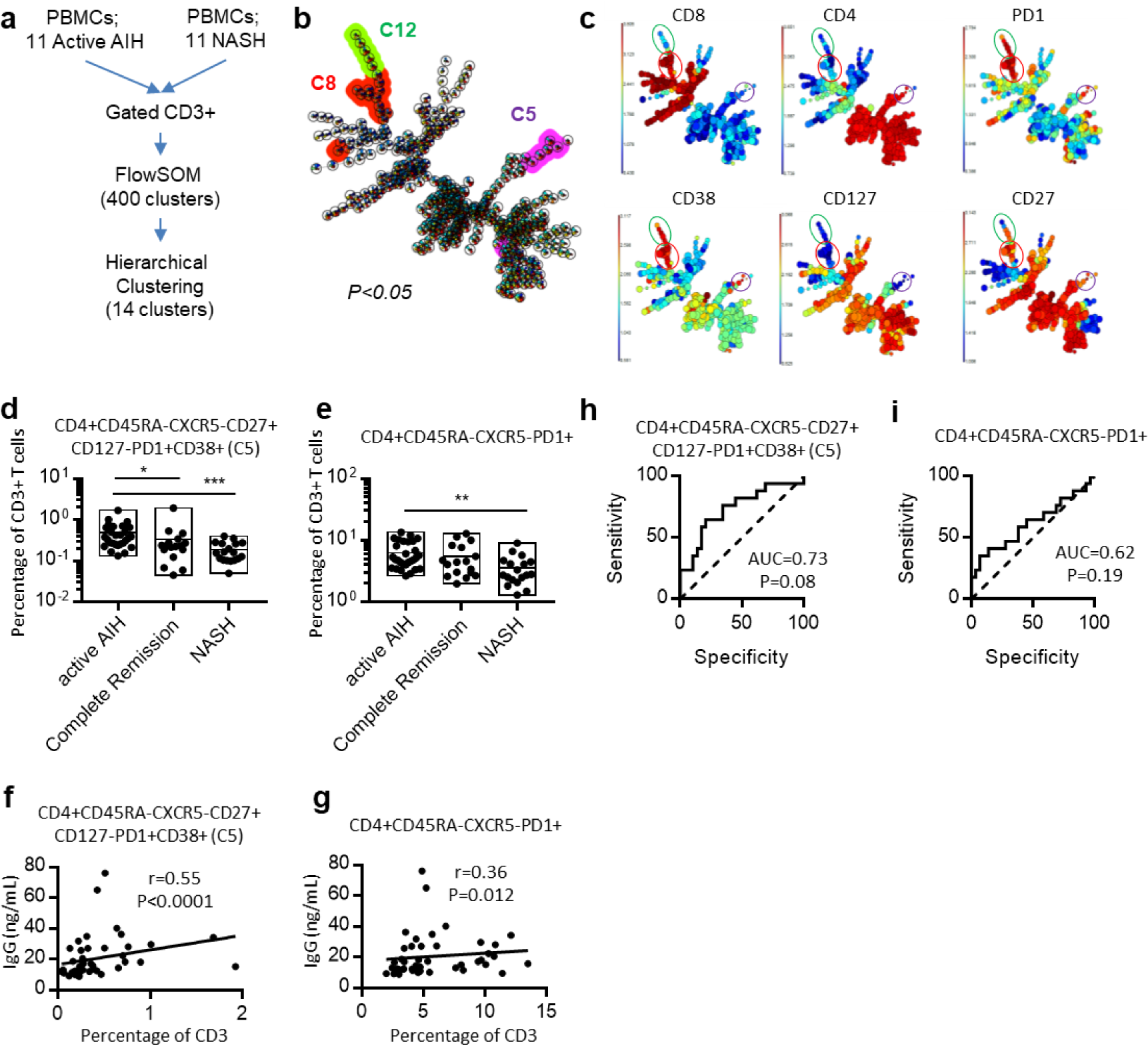
PD-1 and CD38 as potential blood markers of an active AIH. **a**, Strategy of the flow cytometry analysis of the combined T cell population from eleven active AIH patients and eleven NASH patients. **b**, Visualization of the three clusters upregulated in the self-organizing map in AIH patients (p<0,05) after hierarchical clustering of the data generated after FlowSOM. **c**, Visualization of the expression of each marker (indicated in the top of the self-organizing map) within the self-organizing map. Circles indicate the three significant clusters upregulated in AIH patients. **d** and **e**, Graph representation of the percentage of PD-1^+^CD38^+^CXCR5^−^CCR6^−^CD127^−^CD27^+^ (**d**) or PD-1^+^CD38^+^CXCR5^−^ (**e**) memory CD4 T cells within CD3 T cell subset, in active AIH patients (untreated AIH patients (n=19) and AIH patients under treatment but with active disease (n=10)), AIH patients in complete remission under treatment (n=17) and NASH patients (n=18). **f** and **g**, ROC curves analysis between active AIH (n=29) and AIH patients in complete remission under treatment (n=17). **h** and **i**, Spearman correlation between the frequency of CD4 T cell population described in **d** and **e** and the serum IgG level in all AIH patients. Comparisons were performed using the Kruskal-Wallis test and Dunn’s Multiple Comparison Test. *: p<0,05; **: p<0,01; ***: p<0,001.

## Discussion

The combination of the restimulation assay with scRNA-seq, further confirmed by bulk TCRseq, provided strong evidence that the memory PD-1^+^CXCR5^−^CCR6^−^CD27^+^ CD4 T cell population, with B-helper function and a pro-inflammatory signature, is the reservoir of autoreactive CD4 T cells in AIH. This subset supports B cell differentiation through the production of IL-21, which constitutes a new possible therapeutic target for AIH treatment. These results confirm the idea that pathogenic CD4 T cells supporting B cell differentiation in multiple autoimmune disorders share a similar phenotype characterized by the expression of PD-1 and the absence of CXCR5, comparable to the recently described gluten-specific CD4 T cells in celiac disease^18^, TPH cells or TH10 cells in rheumatoid arthritis and in lupus^22–25^. This suggest these cells are not from the classical TFH cell population^18,22–25^. We observed some differences between our study on AIH patients and previous studies on gluten-specific CD4 T cells^18^, autoreactive CD4 T cell described in infants with autoimmune diabetes^20^ and expanded effector CD4 T cells in rheumatoid arthritis^47^. For instance, the PD-1^+^CXCR5^−^ memory CD4 T cell population identified in our study expressed CD27 but did not express the chemokine receptor CCR6.

For the first time we propose a complete transcriptomic analysis of autoreactive (SLA-specific) CD4 T cells in AIH, and we reveal their specific molecular signature characterized by an IL21 and IFNG cytokine signature, resembling TFH (*IL21*) and TPH (*IL21* and *IFNG*) cytokine profiles^22,39,41^. This transcriptomic signature seems to be disease-specific, although similarities with other pathogenic CD4 T cells involved in several autoimmune disorders were observed. Indeed, in autoimmune disorders, distinct pathogenic CD4 T cell molecular signatures have been reported, including a TFH and TPH signature^5,6,9,18,22,24,25^, a TH10 and Tr1/Treg^23,48,49^ (regulatory T cell) signature, and a TH17-like signature^16,19,20,38,50^. Here, we have shown that SLA-specific CD4 T cells had a transcriptomic signature close to the TPH signature^22,24,25^ (*IFNG, IL21, MAF, CD200, ICOS, CTLA4, ITM2A, TIGIT* and *SLAMF6)*. However, *CXCL13*, one of the major genes characterizing TPH cells, was only detected in a small fraction of SLA-specific T cells. SLA-specific CD4 T cells express *IRF4, STAT3, ICOS, CTLA4* and *TIGIT* like TH10 cells in SLE patients^23^ or autoreactive TH17-like cells in MS patients^19^, but their signature was clearly distinct with low expression of *IL10, GZMB, GNLY, IL17, CSF2, IL23A* and *FOSL2.* Finally, we can conclude that SLA-specific CD4 T cells express a specific transcriptomic signature characterized by the transcription factors *NR3C1, EGR2, STAT4, STAT3, IRF4 and MAF;* the cytokines *IL21, IFNG* and *TNF*; and the immune-regulatory molecules *CLEC2D, CTLA4, TIGIT, KLRB1* and *CD109*. These observations demonstrate that SLA-specific CD4 T cells in AIH have a specific molecular profile which partially overlaps with that of recently described TPH cells, and suggest a disease dependent immune signature of autoreactive CD4 T cells. Thus, the direct characterization of the transcriptomic signature of autoreactive CD4 T cells in diverse autoimmune disorders is primordial to elucidate the physiopathology of these cells.

The expression of PD-1 and the overexpression of genes related to immune regulation (*NR3C1, TIGIT* and *CTLA4*) could result from the chronic activation of autoreactive CD4 T cells. However, those autoreactive CD4 T cells persist with a pro-inflammatory profile (*IFNG* and *TNF*), which suggests a form of resistance or adaptation, rather than functional exhaustion. Thus, targeting these pathways (including CD109, a regulator of the TGFβ pathway and of inflammation, or CLEC2D and KLRB1, regulator of the NK killing function) may be interesting new therapeutic options. NK cells are enriched in the liver compared to blood^51^ and a recent study has shown the presence of high numbers of KLRB1^+^ CD4 T cells with a pro-inflammatory cytokine signature (TNF and IFNG) in the inflamed liver ^52^. This suggests that the NK-like signature we observed for autoreactive CD4 T cells in AIH may be imprinted by liver-specific factors.

Finally, based on the data generated from the scRNA-seq and from the phenotype analysis of circulating PD-1^+^CXCR5^−^ CD4 T cells in AIH patients, we have identified a complete specific phenotype of memory CD4 T cells (PD-1^+^CD38^+^CD27^+^CD127^−^CXCR5^−^) associated with an active AIH and linked to serum IgG level. This discovery has immediate clinical relevance for monitoring autoreactive T cells in a small volume of blood during clinical follow-up. The high rate of relapse after immunosuppressive treatment withdrawal is a major challenge when wishing to limit long term immunosuppression and associated side effects in AIH patients. However, classical biological tests (serum transaminases and IgG levels) are not powerful enough to anticipate relapse events. Thus, the capacity to track autoreactive T cell activity in patients’ blood will be of interest for better treatment management in the clinic.

## Acknowledgements

We thank the Bioinformatics Core Facility of Centre d’Immunologie de Marseille-Luminy for helpful discussions and comments. We acknowledge HalioDX and the UCA Genomix platform for sequencing scRNA-seq libraries. This work was supported by institutional grants from INSERM and Nantes Université to the Centre de Recherche en Transplantation et Immunologie, and from INSERM, CNRS and Aix-Marseille University to the Centre d’Immunologie de Marseille-Luminy. This work was carried out in the context of the IHU-Cesti project (ANR-10-IBHU-005) and of the LabEX IGO program (ANR-11-LABX-0016-01), managed by the Agence Nationale de la Recherche. The IHU-Cesti project was also supported by Nantes Métropole and the Région Pays de la Loire. This work was also supported by the patient association ALBI (Association pour la lutte contre les maladies inflammatoires du foie et des voies biliaires), by AFEF-Société Française d’Hépatologie, and by SNFGE-Société Française de Gastro-Entérologie. This work was granted access to the HPC resources of Aix-Marseille Université financed by the project Equip@Meso (ANR-10-EQPX-29-01) of the program “Investissements d’Avenir” supervised by the Agence Nationale de la Recherche.

## References

1. Crotty, S. A brief history of T cell help to B cells. Nat. Rev. Immunol. 15, 185–189 (2015).

2. Fazilleau, N., Mark, L., McHeyzer-Williams, L. J. & McHeyzer-Williams, M. G. Follicular helper T cells: lineage and location. Immunity 30, 324–335 (2009).

3. Morita, R. et al. Human blood CXCR5(+)CD4(+) T cells are counterparts of T follicular cells and contain specific subsets that differentially support antibody secretion. Immunity 34, 108–121 (2011).

4. Yu, D. & Vinuesa C. G. The elusive identity of T follicular helper cells. Trends Immunol. 31, 377–383 (2010).

5. Wang, L. et al. CXCR5+ CD4+ T follicular helper cells participate in the pathogenesis of primary biliary cirrhosis. Hepatology 61, 627–638 (2015).

6. Kenefeck, R. et al. Follicular helper T cell signature in type 1 diabetes. J. Clin. Invest. 125, 292–303 (2015).

7. Kimura, N. et al. Possible involvement of CCR7(-) PD-1(+) follicular helper T cell subset in the pathogenesis of autoimmune hepatitis. J. Gastroenterol. Hepatol. (2017) doi:10.1111/jgh.13844.

8. Ma, L., Qin, J., Ji, H., Zhao, P. & Jiang Y. Tfh and plasma cells are correlated with hypergammaglobulinaemia in patients with autoimmune hepatitis. Liver Int. 34, 405–415 (2014).

9. Weinstein, J. S., Hernandez, S. G. & Craft, J. T cells that promote B-Cell maturation in systemic autoimmunity. Immunol. Rev. 247, 160–171 (2012).

10. Taubert, R., Hupa-Breier, K. L., Jaeckel, E. & Manns M. P. Novel therapeutic targets in autoimmune hepatitis. J. Autoimmun. 95, 34–46 (2018).

11. Sebode, M., Weiler-Normann, C., Liwinski, T. & Schramm, C. Autoantibodies in Autoimmune Liver Disease-Clinical and Diagnostic Relevance. Front Immunol 9, 609 (2018).

12. Sebode, M., Hartl, J., Vergani, D., Lohse, A. W. & International Autoimmune Hepatitis Group (IAIHG). Autoimmune hepatitis: From current knowledge and clinical practice to future research agenda. Liver Int. 38, 15–22 (2018).

13. Renand, A. et al. Immune Alterations in Patients With Type 1 Autoimmune Hepatitis Persist Upon Standard Immunosuppressive Treatment. Hepatol Commun 2, 968–981 (2018).

14. Bovensiepen, C. S. et al. TNF-Producing Th1 Cells Are Selectively Expanded in Liver Infiltrates of Patients with Autoimmune Hepatitis. J. Immunol. (2019) doi:10.4049/jimmunol.1900124.

15. Grant, C. R. et al. Dysfunctional CD39(POS) regulatory T cells and aberrant control of T-helper type 17 cells in autoimmune hepatitis. Hepatology 59, 1007–1015 (2014).

16. Oo, Y. H. et al. CXCR3-dependent recruitment and CCR6-mediated positioning of Th-17 cells in the inflamed liver. J. Hepatol. 57, 1044–1051 (2012).

17. Zhao, L. et al. Interleukin-17 contributes to the pathogenesis of autoimmune hepatitis through inducing hepatic interleukin-6 expression. PLoS ONE 6, e18909 (2011).

18. Christophersen, A. et al. Distinct phenotype of CD4+ T cells driving celiac disease identified in multiple autoimmune conditions. Nat. Med. 25, 734–737 (2019).

19. Cao, Y. et al. Functional inflammatory profiles distinguish myelin-reactive T cells from patients with multiple sclerosis. Sci Transl Med 7, 287ra74 (2015).

20. Heninger, A.-K. et al. A divergent population of autoantigen-responsive CD4+ T cells in infants prior to β cell autoimmunity. Sci Transl Med 9, (2017).

21. Mix, H. et al. Identification of CD4 T-cell epitopes in soluble liver antigen/liver pancreas autoantigen in autoimmune hepatitis. Gastroenterology 135, 2107–2118 (2008).

22. Rao, D. A. et al. Pathologically expanded peripheral T helper cell subset drives B cells in rheumatoid arthritis. Nature 542, 110–114 (2017).

23. Caielli, S. et al. A CD4+ T cell population expanded in lupus blood provides B cell help through interleukin-10 and succinate. Nat. Med. (2018) doi:10.1038/s41591-018-0254-9.

24. Zhang, F. et al. Defining inflammatory cell states in rheumatoid arthritis joint synovial tissues by integrating single-cell transcriptomics and mass cytometry. Nat. Immunol. 20, 928–942 (2019).

25. Arazi, A. et al. The immune cell landscape in kidneys of patients with lupus nephritis. Nat. Immunol. 20, 902–914 (2019).

26. Heneghan, M. A., Yeoman, A. D., Verma, S., Smith, A. D. & Longhi, M. S. Autoimmune hepatitis. Lancet 382, 1433–1444 (2013).

27. Kirstein, M. M. et al. Prediction of short- and long-term outcome in patients with autoimmune hepatitis. Hepatology 62, 1524–1535 (2015).

28. Hennes, E. M. et al. Simplified criteria for the diagnosis of autoimmune hepatitis. Hepatology 48, 169–176 (2008).

29. European Association for the Study of the Liver. EASL Clinical Practice Guidelines: Autoimmune hepatitis. J. Hepatol. 63, 971–1004 (2015).

30. Attaf-Bouabdallah, N. et al. FB5P-seq: FACS-based 5-prime end single-cell RNAseq for integrative analysis of transcriptome and antigen receptor repertoire in B and T cells. http://biorxiv.org/lookup/doi/10.1101/795575 (2019) doi:10.1101/795575.

31. Macosko, E. Z. et al. Highly Parallel Genome-wide Expression Profiling of Individual Cells Using Nanoliter Droplets. Cell 161, 1202–1214 (2015).

32. Stuart, T. et al. Comprehensive Integration of Single-Cell Data. Cell 177, 1888–1902.e21 (2019).

33. Alamyar, E., Duroux, P., Lefranc, M.-P. & Giudicelli, V. IMGT(®) tools for the nucleotide analysis of immunoglobulin (IG) and T cell receptor (TR) V-(D)-J repertoires, polymorphisms, and IG mutations: IMGT/V-QUEST and IMGT/HighV-QUEST for NGS. Methods Mol. Biol. 882, 569–604 (2012).

34. Renand, A. et al. Chronic cat allergen exposure induces a TH2 cell-dependent IgG4 response related to low sensitization. J. Allergy Clin. Immunol. (2015) doi:10.1016/j.jaci.2015.07.031.

35. Renand, A. et al. Heterogeneity of Ara h Component-Specific CD4 T Cell Responses in Peanut-Allergic Subjects. Front Immunol 9, 1408 (2018).

36. Bacher, P. et al. Antigen-reactive T cell enrichment for direct, high-resolution analysis of the human naive and memory Th cell repertoire. J. Immunol. 190, 3967–3976 (2013).

37. Conti, H. R. & Gaffen, S. L. IL-17-Mediated Immunity to the Opportunistic Fungal Pathogen Candida albicans. J. Immunol. 195, 780–788 (2015).

38. Zhu, J., Yamane, H. & Paul, W. E. Differentiation of effector CD4 T cell populations (*). Annu. Rev. Immunol. 28, 445–489 (2010).

39. Weinstein, J. S. et al. Global transcriptome analysis and enhancer landscape of human primary T follicular helper and T effector lymphocytes. Blood 124, 3719–3729 (2014).

40. Ciofani, M. et al. A validated regulatory network for Th17 cell specification. Cell 151, 289–303 (2012).

41. Locci, M. et al. Human circulating PD-1+CXCR3-CXCR5+ memory Tfh cells are highly functional and correlate with broadly neutralizing HIV antibody responses. Immunity 39, 758–769 (2013).

42. Cain, D. W. & Cidlowski, J. A. Immune regulation by glucocorticoids. Nat. Rev. Immunol. 17, 233–247 (2017).

43. Litvinov, I. V. et al. CD109 release from the cell surface in human keratinocytes regulates TGF-β receptor expression, TGF-β signalling and STAT3 activation: relevance to psoriasis. Exp. Dermatol. 20, 627–632 (2011).

44. Tsai, Y.-L. et al. Endoplasmic reticulum stress activates SRC, relocating chaperones to the cell surface where GRP78/CD109 blocks TGF-β signaling. Proc. Natl. Acad. Sci. U.S.A. 115, E4245–E4254 (2018).

45. Song, G. et al. CD109 regulates the inflammatory response and is required for the pathogenesis of rheumatoid arthritis. Ann. Rheum. Dis. (2019) doi:10.1136/annrheumdis-2019-215473.

46. Llibre, A., Klenerman, P. & Willberg, C. B. Multi-functional lectin-like transcript-1: A new player in human immune regulation. Immunol. Lett. 177, 62–69 (2016).

47. Fonseka, C. Y. et al. Mixed-effects association of single cells identifies an expanded effector CD4+ T cell subset in rheumatoid arthritis. Sci Transl Med 10, (2018).

48. Buckner, J. H. Mechanisms of impaired regulation by CD4(+)CD25(+)FOXP3(+) regulatory T cells in human autoimmune diseases. Nat. Rev. Immunol. 10, 849–859 (2010).

49. Roncarolo, M. G., Gregori, S., Bacchetta, R., Battaglia, M. & Gagliani, N. The Biology of T Regulatory Type 1 Cells and Their Therapeutic Application in Immune-Mediated Diseases. Immunity 49, 1004–1019 (2018).

50. Paroni, M. et al. Recognition of viral and self-antigens by TH1 and TH1/TH17 central memory cells in patients with multiple sclerosis reveals distinct roles in immune surveillance and relapses. J. Allergy Clin. Immunol. (2017) doi:10.1016/j.jaci.2016.11.045.

51. Heymann, F. & Tacke, F. Immunology in the liver--from homeostasis to disease. Nat Rev Gastroenterol Hepatol 13, 88–110 (2016).

52. Truong, K.-L. et al. Killer-like receptors and GPR56 progressive expression defines cytokine production of human CD4+ memory T cells. Nat Commun 10, 2263 (2019).

