## Extended Data Figure 1 for "Integrative molecular profiling of autoreactive CD4 T cells in autoimmune hepatitis"

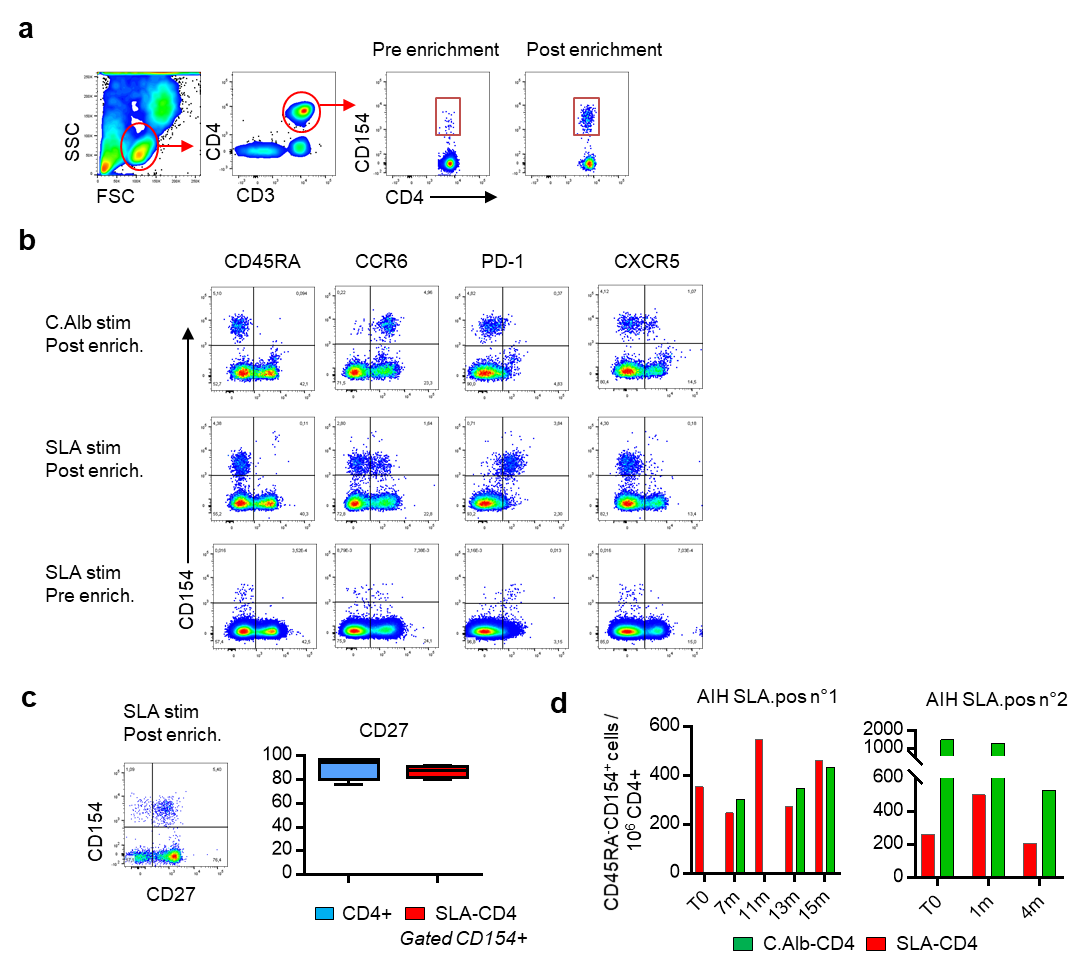

**Extended Data Figure 1. Detection of CD154^+^CD4^+^ T cells after peptide stimulation assay.** **a**, Gating strategy to detect CD154^+^CD4^+^ T cells after an enrichment step on magnetic columns. **b**, Dot plot representation on gated CD3^+^CD4^+^ T cells of CD154 expression versus markers indicated above, before or after enrichment. **c**, Dot plot representation on gated CD3^+^CD4^+^ T cells of CD154 expression versus CD27. **d**, Longitudinal analysis of the frequency of CD154^+^CD4^+^CD45RA^-^ cells per million total CD4 T cells from two distinct AIH SLA-pos patients.
