## Extended Data Figure 2 for "Integrative molecular profiling of autoreactive CD4 T cells in autoimmune hepatitis"

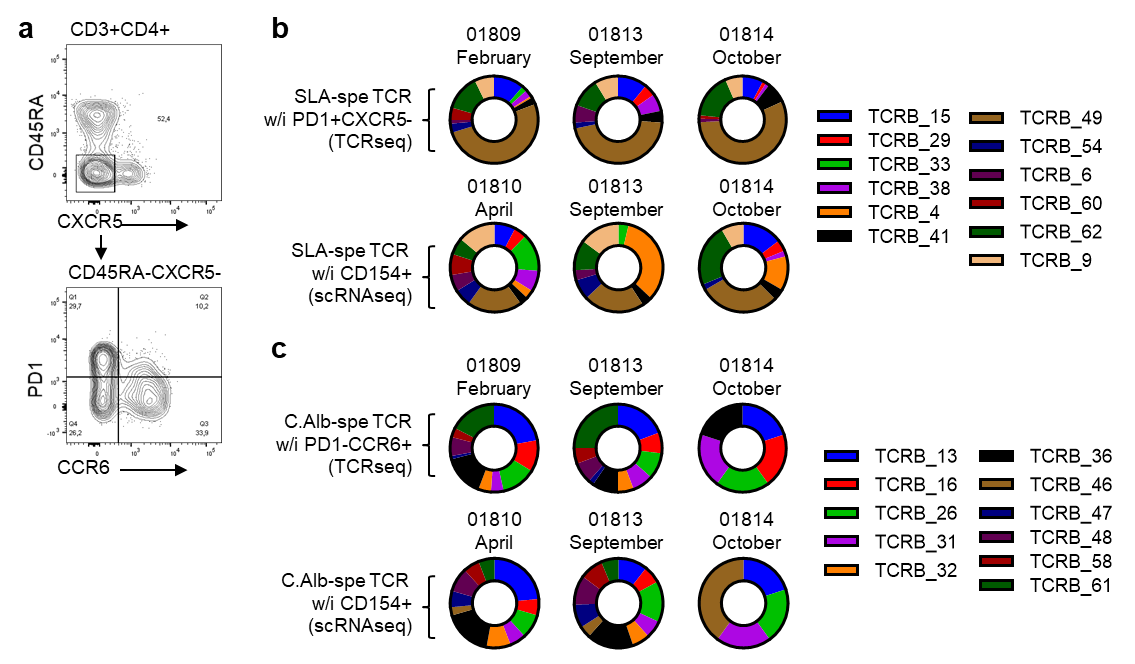

**Extended Data Figure 2. Stable specific-TCR distribution over time.** **a**, Gating strategy to isolate PD-1^+^CXCR5^-^CCR6^-^, PD-1^-^CCR6^+^CXCR5^-^ and PD-1^-^CXCR5^-^CCR6^-^ CD4 memory T cells in the blood of patient 018. **b** and **c**, Quantitative distribution of the identified TCRβ of SLA-CD4 T cells (**b**) or of C.Alb-CD4 T cells (**c**) at three different time point and from scRNAseq data analysis.
