## Extended Data Figure 3 for "Integrative molecular profiling of autoreactive CD4 T cells in autoimmune hepatitis"

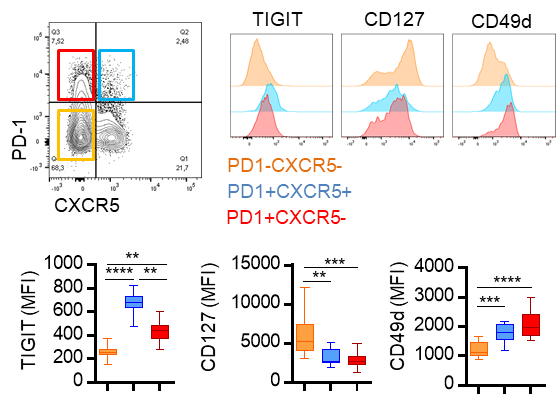

**Extended Data Figure 3. Phenotype of the PD-1^+^CXCR5^-^ CD4 T cell population in the peripheral blood of AIH patients.** PD-1 and CXCR5 expression by memory CD4^+^ T cells in one AIH patient. Median of fluorescence intensity for each marker indicated in PD-1^+^CXCR5^-^ (red), PD-1^+^CXCR5^+^ (blue) and PD-1^-^CXCR5^-^ (orange) memory CD4 T cell subsets in AIH patients (n=12). Comparisons were performed using the Kruskal-Wallis test and Dunn's Multiple Comparison Test. *: p<0,05; **: p<0,01; ***: p<0,001; ****: p<0,0001.
