## Extended Data Figure 4 for "Integrative molecular profiling of autoreactive CD4 T cells in autoimmune hepatitis"

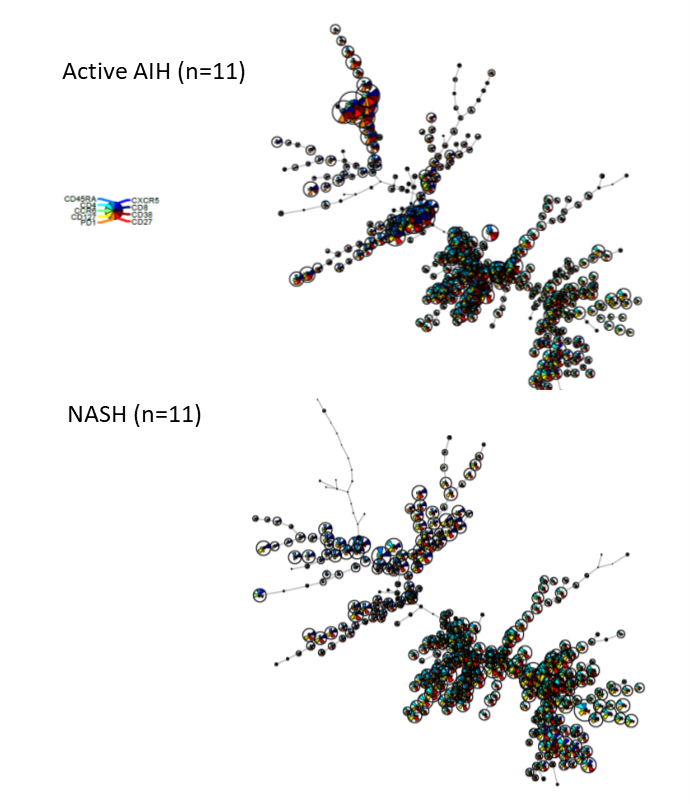

**Extended Data Figure 4. FlowSOM representation of flow cytometry data.** Self-organizing maps by using FlowSOM (400 clusters) of the CD3 T cell populations from active AIH patients (top, n=11) and NASH patients (bottom, n=11). Each points represent a cluster of cell sharing similar profile of expression.
