## Extended Data Figure 5 for "Integrative molecular profiling of autoreactive CD4 T cells in autoimmune hepatitis"

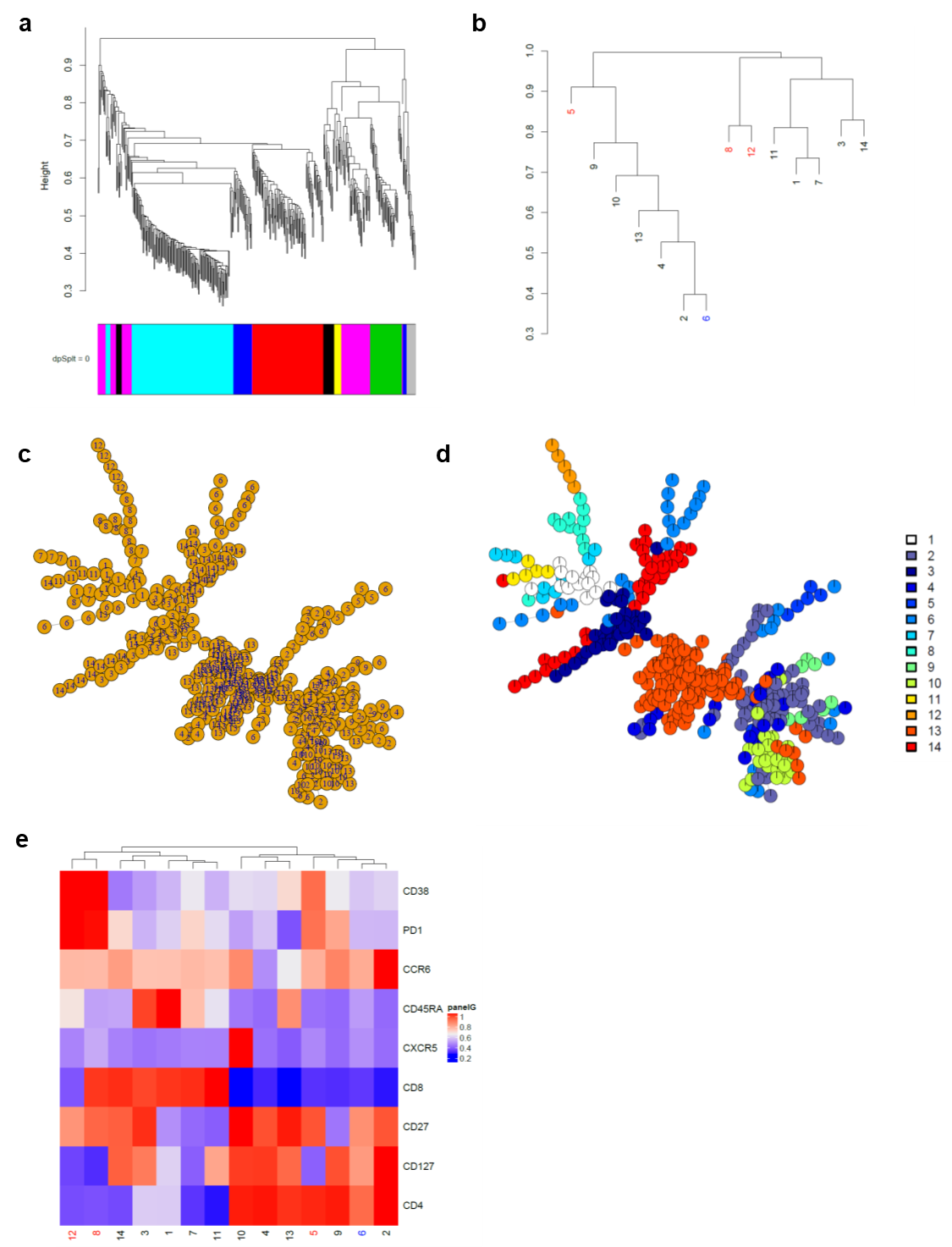

**Extended Data Figure 5. Metaclusterization of the data generated using flow cytometry data and FlowSOM approach.** **a**, Hierarchical clustering of the data generated after FlowSOM. **b**, Hierarchical clustering with each 14 clusters indicated. In red, cellular clusters significantly upregulated in AIH patients. In blue, cellular cluster significantly upregulated in NASH patients. **c** and **d**, The 14 clusters were identify in the self-organizing maps by numbers (**c**) or colors (**d**). **e**, Heat-map of each marker for the 14 clusters identified in the total CD3 population (AIH and NASH).
