## Extended Data Figure 6 for "Integrative molecular profiling of autoreactive CD4 T cells in autoimmune hepatitis"

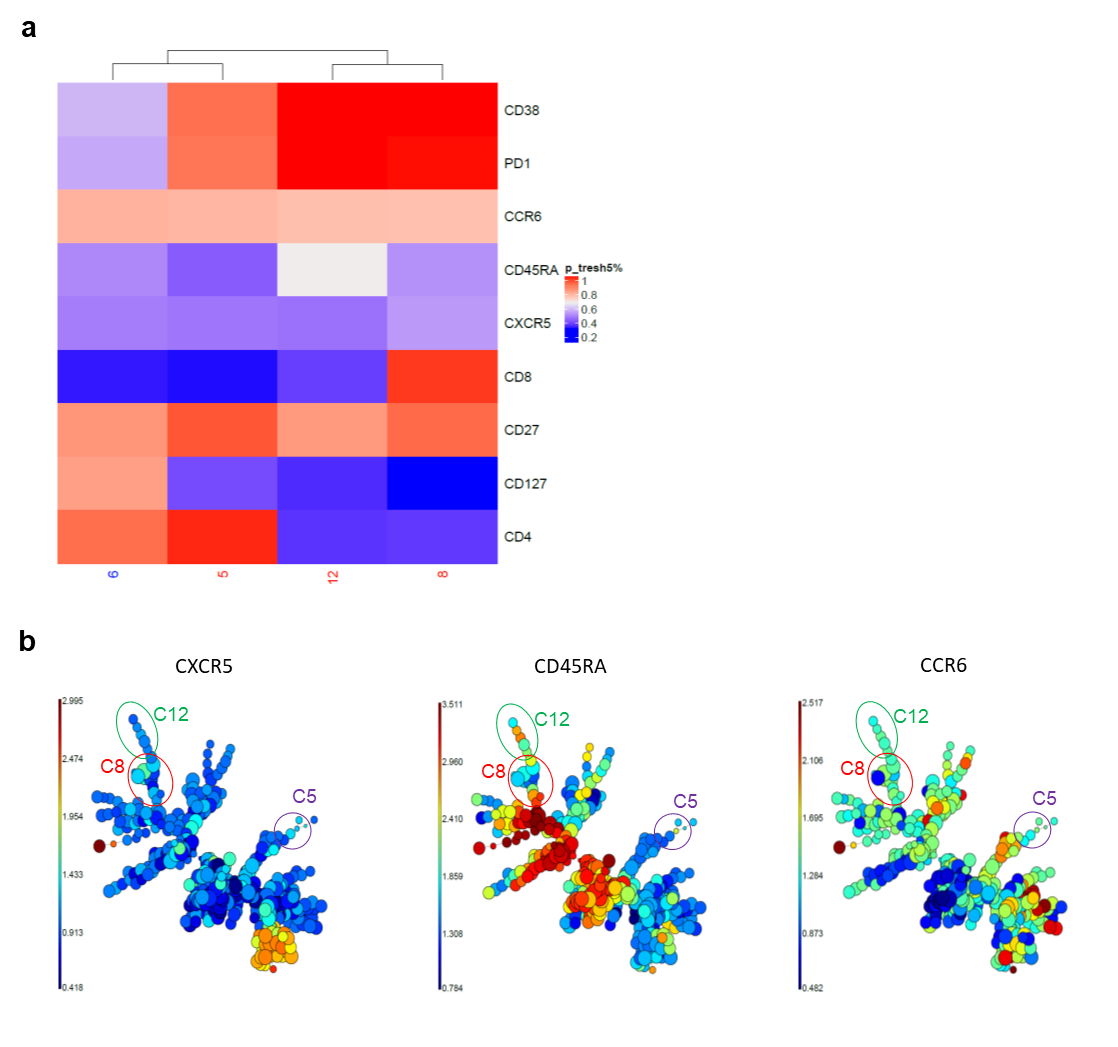

**Extended Data Figure 6. Characterization of T cell clusters significantly modulated in AIH patients.** **a**, Heat-map representation of each marker for up regulated clusters in AIH patients (red) or in NASH patients (blue) with a p value < 0.05. **b**, Visualization of the expression of each marker (indicated in the top of the self-organizing map) within the self-organizing map. Circles indicate the three significant clusters upregulated in AIH patients.
