## Extended Data Figure 7 for "Integrative molecular profiling of autoreactive CD4 T cells in autoimmune hepatitis"

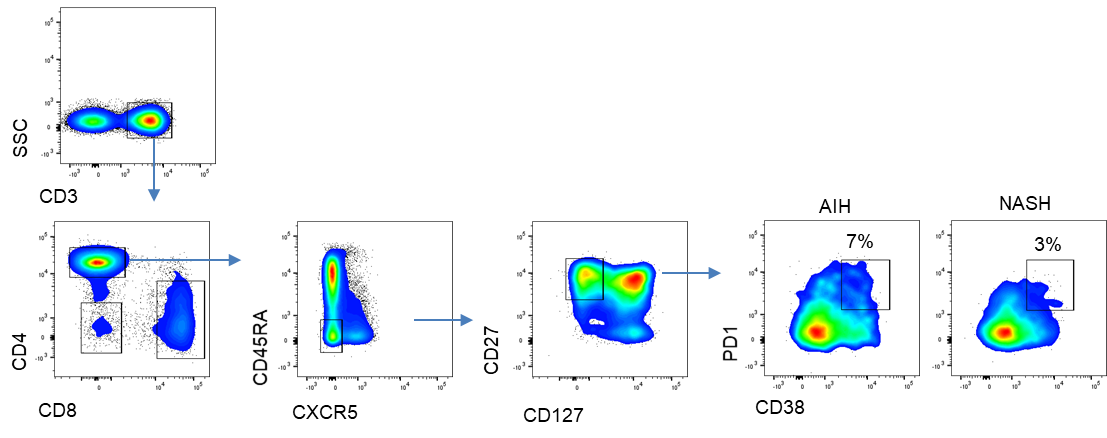

**Extended Data Figure 7. Supervised analysis by flow cytometry of the PD1 and CD38 expression at the surface of CD4 T cell subset.** Flow cytometry gating strategy to measure PD1 and CD38 expression at the surface of CD4 T cell subset identify after metaclustering analysis.
