## Extended Table 1 for "Integrative molecular profiling of autoreactive CD4 T cells in autoimmune hepatitis"

| **Batch** | **Samples** | **# plates** | **# cells sorted** | **# cells passing QC (%)** | **Mean sequencing depth (on cells passing QC)** | **Mean gene detection** | **# cells with TCRα and TCRβ sequence (%)** | **# cells with only TCRβ sequence (%)** | **# cells with only TCRα sequence (%)** | **# cells with no TCR sequence (%)** |
| --- | --- | --- | --- | --- | --- | --- | --- | --- | --- | --- |
| 180523 | 018-10, 028-03, 128-01 | 7 | 545 | 517 (95%) | 439,911 reads/cell | 972 genes/cell | 185 (36%) | 182 (35%) | 43 (8%) | 107 (21%) |
| 190220 | 004-05, 018-13, 018-14 | 6 | 432 | 382 (88%) | 428,664 reads/cell | 1426 genes/cell | 161 (42%) | 94 (25%) | 32 (8%) | 95 (25%) |
| 190612 | 051-04 | 2 | 146 | 140 (96%) | 332,635 reads/cell | 517 genes/cell | 23 (16%) | 32 (23%) | 21 (15%) | 64 (46%) |

**Extended Table 1. scRNAseq data: metrics and quality controls**
