## Extended Table 2 for "Integrative molecular profiling of autoreactive CD4 T cells in autoimmune hepatitis"

| **gene** | **cluster** | **avg_logFC** | **pct.1** | **pct.2** | **diff.pct** | **p_val_adj** |
| --- | --- | --- | --- | --- | --- | --- |
| IL22 | Calb | 3.7812791394249 | 0.935 | 0.328 | 0.607 | 7.04619348480492e-182 |
| IL2 | Calb | 2.89948572176832 | 0.901 | 0.394 | 0.507 | 1.96135596918873e-140 |
| CCL20 | Calb | 2.79548155105928 | 0.759 | 0.24 | 0.519 | 1.10240821657001e-130 |
| CSF2 | Calb | 3.09806871956395 | 0.767 | 0.256 | 0.511 | 1.49414965400622e-118 |
| IL17A | Calb | 3.01748818790306 | 0.59 | 0.114 | 0.476 | 9.67409597290935e-106 |
| TNFSF14 | Calb | 1.46682145062625 | 0.73 | 0.3 | 0.43 | 9.22435278510163e-77 |
| MYC | Calb | 1.38565789930743 | 0.751 | 0.264 | 0.487 | 3.15128845092905e-64 |
| AC020571.3 | Calb | 1.93960967780003 | 0.586 | 0.143 | 0.443 | 5.99601264196115e-62 |
| TNF | Calb | 1.20604683951415 | 0.874 | 0.612 | 0.262 | 7.33205042713228e-54 |
| GPR171 | Calb | 1.52924572806296 | 0.643 | 0.233 | 0.41 | 2.19071102392586e-52 |
| IL23A | Calb | 1.77454679389244 | 0.55 | 0.159 | 0.391 | 3.79634233303616e-48 |
| LTA | Calb | 1.87304244363393 | 0.471 | 0.099 | 0.372 | 3.31731645930913e-47 |
| FASLG | Calb | 1.40140901432786 | 0.475 | 0.09 | 0.385 | 5.31287779373244e-45 |
| KLF10 | Calb | 1.40379349287521 | 0.552 | 0.165 | 0.387 | 1.36605093041026e-44 |
| DUSP2 | Calb | 1.18740530606693 | 0.623 | 0.386 | 0.237 | 2.20038176397292e-33 |
| PHLDA1 | Calb | 1.16397535097906 | 0.686 | 0.364 | 0.322 | 2.31762137920636e-33 |
| VIM | Calb | 1.02463408827461 | 0.507 | 0.156 | 0.351 | 4.01441648115644e-31 |
| HMGA1 | Calb | 0.877977618466401 | 0.469 | 0.132 | 0.337 | 2.75583439955628e-29 |
| CD48 | Calb | 1.1005745255796 | 0.57 | 0.222 | 0.348 | 8.57695509289518e-29 |
| NOP16 | Calb | 0.866011318775976 | 0.68 | 0.337 | 0.343 | 1.49064800164169e-28 |
| NINJ1 | Calb | 1.37111857210451 | 0.408 | 0.128 | 0.28 | 3.84138230256711e-28 |
| FOSL1 | Calb | 1.14160568379372 | 0.373 | 0.081 | 0.292 | 4.23568673374345e-28 |
| GADD45B | Calb | 0.920043190264482 | 0.783 | 0.529 | 0.254 | 4.36840219478028e-28 |
| PTGER2 | Calb | 1.39720063853642 | 0.266 | 0.024 | 0.242 | 2.57804019506894e-27 |
| REL | Calb | 1.02531479474298 | 0.663 | 0.344 | 0.319 | 5.04310887175995e-26 |
| SATB1 | Calb | 1.07837019064749 | 0.485 | 0.185 | 0.3 | 1.03790702896669e-24 |
| SPRY1 | Calb | 0.880903711824148 | 0.519 | 0.196 | 0.323 | 1.94635563848823e-24 |
| NR4A3 | Calb | 0.752451975667876 | 0.787 | 0.557 | 0.23 | 8.40914053055642e-24 |
| DDX21 | Calb | 0.761522579631821 | 0.677 | 0.383 | 0.294 | 2.77698647509408e-23 |
| IL7R | Calb | 0.668501913957269 | 0.781 | 0.527 | 0.254 | 5.88424769270102e-20 |
| GPR183 | Calb | 1.01085829824314 | 0.428 | 0.181 | 0.247 | 7.50772933690466e-19 |
| PKIA | Calb | 0.573140534764982 | 0.383 | 0.115 | 0.268 | 2.15409572471559e-18 |
| BMI1 | Calb | 0.554553048954698 | 0.312 | 0.079 | 0.233 | 2.17155394813178e-16 |
| RASSF5 | Calb | 0.692815236218909 | 0.706 | 0.452 | 0.254 | 7.66947435188068e-16 |
| WDR43 | Calb | 0.759501078779376 | 0.511 | 0.234 | 0.277 | 8.44576165800225e-16 |
| NXT1 | Calb | 0.773657088079381 | 0.387 | 0.158 | 0.229 | 1.03512533375503e-15 |
| BACH2 | Calb | 0.828851592422762 | 0.426 | 0.201 | 0.225 | 1.52438674853158e-14 |
| TNFRSF4 | Calb | 0.904919971026685 | 0.509 | 0.269 | 0.24 | 6.20466791712204e-14 |
| RUNX3 | Calb | 0.402786043621213 | 0.264 | 0.062 | 0.202 | 1.53175048703788e-13 |
| IFRD2 | Calb | 0.577116999947271 | 0.276 | 0.071 | 0.205 | 1.57397184514884e-13 |
| FOSL2 | Calb | 0.609853108701106 | 0.424 | 0.187 | 0.237 | 3.42104351976174e-13 |
| KDM6B | Calb | 0.713549581646057 | 0.574 | 0.319 | 0.255 | 1.38810631335598e-12 |
| PIM3 | Calb | 0.584390447831632 | 0.501 | 0.253 | 0.248 | 1.91145263574056e-12 |
| CYCS | Calb | 0.59951583302444 | 0.744 | 0.524 | 0.22 | 2.29813376054965e-11 |
| RGCC | Calb | 0.573247158442548 | 0.726 | 0.516 | 0.21 | 4.01298778960846e-11 |
| EIF5A | Calb | 0.622847080062699 | 0.493 | 0.267 | 0.226 | 3.34965779958374e-09 |
| BTG1 | Calb | 0.525092002623129 | 0.694 | 0.48 | 0.214 | 1.26936942707063e-08 |
| TFRC | Calb | 0.617382900581562 | 0.391 | 0.189 | 0.202 | 1.10918859699876e-07 |
| RPS2 | Calb | 0.462089079417107 | 0.497 | 0.282 | 0.215 | 5.30980835828458e-07 |
| KMT2E-AS1 | Calb | 0.33504451932403 | 0.473 | 0.266 | 0.207 | 1.99149195446352e-06 |
| NR3C1 | SLA | 1.50464099817351 | 0.767 | 0.32 | 0.447 | 1.77584594191694e-60 |
| CD109 | SLA | 1.41487573722481 | 0.5 | 0.075 | 0.425 | 3.04117374026698e-50 |
| IGFL2 | SLA | 1.97245115146088 | 0.311 | 0.014 | 0.297 | 7.36822309599279e-45 |
| IL21 | SLA | 1.35545256262815 | 0.678 | 0.262 | 0.416 | 2.21623845509595e-44 |
| CLEC2D | SLA | 0.613978581706778 | 0.894 | 0.59 | 0.304 | 2.12923454863985e-42 |
| CTLA4 | SLA | 1.13206319557209 | 0.582 | 0.178 | 0.404 | 2.5451217791417e-40 |
| TIGIT | SLA | 1.59624980672014 | 0.419 | 0.065 | 0.354 | 1.2313684847282e-39 |
| ITM2A | SLA | 1.00371061385282 | 0.745 | 0.398 | 0.347 | 5.67501044331078e-36 |
| TBC1D4 | SLA | 1.24087322419055 | 0.535 | 0.178 | 0.357 | 1.31390632190881e-33 |
| MAGEH1 | SLA | 1.66134573005475 | 0.3 | 0.043 | 0.257 | 5.45305871608269e-30 |
| ITGA4 | SLA | 1.1981745309445 | 0.368 | 0.069 | 0.299 | 2.59865373305932e-28 |
| RDH10 | SLA | 1.05536848548306 | 0.297 | 0.03 | 0.267 | 2.78666615036777e-28 |
| IFITM1 | SLA | 0.744724529846605 | 0.806 | 0.564 | 0.242 | 4.72944063100852e-27 |
| GK | SLA | 0.86405861649931 | 0.495 | 0.158 | 0.337 | 1.27565572786878e-26 |
| ARHGAP26 | SLA | 1.58783095084319 | 0.355 | 0.083 | 0.272 | 1.18102915915552e-25 |
| IFNG | SLA | 0.80594047764301 | 0.551 | 0.215 | 0.336 | 3.61333302220941e-25 |
| LYST | SLA | 0.807846786084541 | 0.59 | 0.3 | 0.29 | 8.2482159755313e-22 |
| MAF | SLA | 0.846205644345728 | 0.31 | 0.059 | 0.251 | 5.25840963770986e-21 |
| ZEB2 | SLA | 0.934600076032697 | 0.302 | 0.055 | 0.247 | 7.9047498416679e-21 |
| AIM1 | SLA | 1.10603400880223 | 0.516 | 0.264 | 0.252 | 1.58832188044701e-20 |
| SGK1 | SLA | 1.28227453105417 | 0.379 | 0.114 | 0.265 | 3.28908467890336e-20 |
| NDUFV2P1 | SLA | 0.292714345209839 | 0.441 | 0.154 | 0.287 | 7.28485590972116e-20 |
| NIN | SLA | 0.903715602942602 | 0.43 | 0.144 | 0.286 | 1.01444923820973e-19 |
| H2AFZ | SLA | 0.702961006584069 | 0.749 | 0.538 | 0.211 | 6.22277056853403e-19 |
| PSMB9 | SLA | 1.09452281139659 | 0.332 | 0.089 | 0.243 | 5.49289574518635e-17 |
| MPHOSPH8 | SLA | 0.923051171652394 | 0.44 | 0.176 | 0.264 | 6.00818283262027e-17 |
| FKBP5 | SLA | 0.968221718498138 | 0.388 | 0.136 | 0.252 | 6.44095926295351e-17 |
| SLAMF6 | SLA | 1.03794666004249 | 0.319 | 0.089 | 0.23 | 8.08597006916759e-17 |
| SOD1 | SLA | 0.490566520165665 | 0.874 | 0.677 | 0.197 | 1.06988564809042e-16 |
| AC006129.2 | SLA | 0.864246166823883 | 0.364 | 0.112 | 0.252 | 1.82807744818666e-16 |
| GCC2 | SLA | 0.98888387854577 | 0.337 | 0.099 | 0.238 | 1.29706523591118e-15 |
| FABP5 | SLA | 0.746563686174373 | 0.509 | 0.272 | 0.237 | 7.66080441653487e-14 |
| HNRNPLL | SLA | 0.979629450896608 | 0.401 | 0.172 | 0.229 | 8.39022745771018e-14 |
| MIR4435-2HG | SLA | 0.531784911055858 | 0.522 | 0.264 | 0.258 | 1.56820902430298e-12 |
| CIRBP | SLA | 0.899121212329784 | 0.46 | 0.245 | 0.215 | 2.71420741728082e-12 |
| KLRB1 | SLA | 0.664987401620866 | 0.505 | 0.292 | 0.213 | 5.539908321737e-12 |
| SYNE2 | SLA | 0.875530288263169 | 0.399 | 0.178 | 0.221 | 1.08668785527918e-10 |
| MAPRE2 | SLA | 0.672866865458566 | 0.614 | 0.371 | 0.243 | 2.21993246451479e-10 |
| DYNLL1 | SLA | 0.944016988522934 | 0.359 | 0.152 | 0.207 | 2.49054322157501e-10 |
| YPEL5 | SLA | 0.864389192046447 | 0.397 | 0.183 | 0.214 | 1.27178120462173e-09 |
| CD59 | SLA | 0.851249913043773 | 0.412 | 0.201 | 0.211 | 1.6453758451645e-09 |
| ANXA5 | SLA | 0.851074002620189 | 0.364 | 0.16 | 0.204 | 3.62309538871449e-09 |
| SLC20A1 | SLA | 0.526998302993381 | 0.397 | 0.191 | 0.206 | 2.02947677396629e-08 |
| GBP2 | SLA | 0.558329070459029 | 0.687 | 0.481 | 0.206 | 4.89100659836978e-08 |
| CDC42SE2 | SLA | 0.557972211759629 | 0.438 | 0.223 | 0.215 | 5.35373991440258e-08 |
| RAB27A | SLA | 0.766690131496838 | 0.454 | 0.241 | 0.213 | 6.90410825086938e-08 |
| AHI1 | SLA | 0.639331857846134 | 0.557 | 0.351 | 0.206 | 9.24334029157627e-07 |
| HMGN2 | SLA | 0.6166361375444 | 0.441 | 0.237 | 0.204 | 1.68585985466152e-06 |
| CAP1 | SLA | 0.593886037841836 | 0.452 | 0.256 | 0.196 | 1.96938994324569e-06 |
| TNFAIP8 | SLA | 0.507307216569338 | 0.586 | 0.377 | 0.209 | 5.70593317025512e-06 |

**Extended Table 2. Top 50 significantly differentially expressed genes between C.Alb-specific and SLA-specific CD4 T cells**
