## Extended Table 3 for "Integrative molecular profiling of autoreactive CD4 T cells in autoimmune hepatitis"

|  |  |  |  |  |  | **TCRA** | | | | **TCRB** | | |
| --- | --- | --- | --- | --- | --- | --- | --- | --- | --- | --- | --- | --- |
| **TCR name** | **Specificity** | **Patient ID** | **nbr clones** | **TCRA** | **TCRB** | **V-GENE and allele** | **J-GENE and allele** | **AA JUNCTION** | **V-GENE and allele** | | **J-GENE and allele** | **AA JUNCTION** |
| a1_b60 | SLA | #018 | 5 | TCRA_1 | TCRB_60 | Homsap TRAV21*02 (F) | Homsap TRAJ48*01 F | CAAPAAGNEKLTF | Homsap TRBV20-1*02 F | | Homsap TRBJ2-3*01 F | CSAGDRTGDTDTQYF |
| a10_b62 | SLA | #018 | 18 | TCRA_10 | TCRB_62 | Homsap TRAV19*01 F | Homsap TRAJ42*01 F | CALSEAGGSQGNLIF | Homsap TRBV20-1*02 F | | Homsap TRBJ1-1*01 F | CSASRGQGLVNTEAFF |
| a13_b6 | SLA | #018 | 5 | TCRA_13 | TCRB_6 | Homsap TRAV19*01 F | Homsap TRAJ5*01 F | CALSQPLTGRRALTF | Homsap TRBV7-2*01 F | | Homsap TRBJ2-4*01 F | CASSFEGATDIQYF |
| a14_b29 | SLA | #018 | 5 | TCRA_14 | TCRB_29 | Homsap TRAV9-2*01 F | Homsap TRAJ42*01 F | CALSTGYGGSQGNLIF | Homsap TRBV7-2*01 F | | Homsap TRBJ2-1*01 F | CASSLQLAAINEQFF |
| a15 | SLA | #018 | 2 | TCRA_15 | #N/A | Homsap TRAV9-2*03 (F) | Homsap TRAJ43*01 F | CALSVQMYNNNDMRF | #N/A | | #N/A | #N/A |
| a16_b33 | SLA | #018 | 10 | TCRA_16 | TCRB_33 | Homsap TRAV21*02 (F) | Homsap TRAJ11*01 F | CAMSQDRNSGYSTLTF | Homsap TRBV18*01 F | | Homsap TRBJ2-3*01 F | CASSPAGAADTQYF |
| a17_b23 | SLA | #018 | 3 | TCRA_17 | TCRB_23 | Homsap TRAV12-3*01 F | Homsap TRAJ54*01 F | CAMSVPQGAQKLVF | Homsap TRBV7-2*01 F | | Homsap TRBJ2-3*01 F | CASSLGGGTDTQYF |
| a2_b44 | SLA | #018 | 2 | TCRA_2 | TCRB_44 | Homsap TRAV29/DV5*01 F | Homsap TRAJ57*01 F | CAASSQGGSEKLVF | Homsap TRBV4-1*01 F | | Homsap TRBJ2-6*01 F | CASSQDPIGGAAGANVLTF |
| a23_b5 | SLA | #018 | 2 | TCRA_23 | TCRB_5 | Homsap TRAV2*01 F | Homsap TRAJ29*01 F | CAVEDRSGNTPLVF | Homsap TRBV7-2*01 F | | Homsap TRBJ2-5*01 F | CASSEQRGAQETQYF |
| a24_b53 | SLA | #018 | 4 | TCRA_24 | TCRB_53 | Homsap TRAV22*01 F | Homsap TRAJ11*01 F | CAVESSGYSTLTF | Homsap TRBV7-2*01 F | | Homsap TRBJ2-3*01 F | CASSTQGTDTQYF |
| a26_b21 | SLA | #018 | 3 | TCRA_26 | TCRB_21 | Homsap TRAV2*01 F | Homsap TRAJ37*01 F | CAVGPGNTGKLIF | Homsap TRBV7-2*01 F | | Homsap TRBJ2-3*01 F | CASSLESSDTQYF |
| a27_b9 | SLA | #018 | 6 | TCRA_27 | TCRB_9 | Homsap TRAV8-3*02 (F) | Homsap TRAJ4*01 F | CAVGSGGYNKLIF | Homsap TRBV7-2*02 F, or Homsap TRBV7-2*03 F | | Homsap TRBJ1-2*01 F | CASSFPGQGVYGYTF |
| a28_b54 | SLA | #018 | 7 | TCRA_28 | TCRB_54 | Homsap TRAV8-3*02 (F) | Homsap TRAJ10*01 F | CAVGVTGGGNKLTF | Homsap TRBV12-4*01 F | | Homsap TRBJ1-6*02 F | CASSTSRTGSPLHF |
| a29_b57 | SLA | #018 | 4 | TCRA_29 | TCRB_57 | Homsap TRAV8-1*01 F | Homsap TRAJ29*01 F | CAVNAPNSGNTPLVF | Homsap TRBV7-2*01 F | | Homsap TRBJ2-3*01 F | CASSVGLAGAAGADTQYF |
| a30_b35 | SLA | #018 | 2 | TCRA_30 | TCRB_35 | Homsap TRAV21*02 (F) | Homsap TRAJ40*01 F | CAVPVGTYKYIF | Homsap TRBV12-3*01 F | | Homsap TRBJ1-3*01 F | CASSPPEGRGNTIYF |
| a31_b14 | SLA | #018 | 2 | TCRA_31 | TCRB_14 | Homsap TRAV20*02 (F) | Homsap TRAJ33*01 F | CAVQRDSNYQLIW | Homsap TRBV3-1*01 F | | Homsap TRBJ1-2*01 F | CASSHWDRGDYGYTF |
| a32_b9 | SLA | #018 | 11 | TCRA_32 | TCRB_9 | Homsap TRAV3*01 F | Homsap TRAJ13*01 F | CAVRDIGGGYQKVTF | Homsap TRBV7-2*02 F, or Homsap TRBV7-2*03 F | | Homsap TRBJ1-2*01 F | CASSFPGQGVYGYTF |
| a35 | SLA | #018 | 2 | TCRA_35 | #N/A | Homsap TRAV41*01 F | Homsap TRAJ49*01 F | CAVSDGNQFYF | #N/A | | #N/A | #N/A |
| a38_b49 | SLA | #018 | 15 | TCRA_38 | TCRB_49 | Homsap TRAV8-1*01 F | Homsap TRAJ13*01 F | CAVTPSGGYQKVTF | Homsap TRBV7-3*01 F | | Homsap TRBJ1-2*01 F | CASSSFDRGNYGYTF |
| a39_b15 | SLA | #018 | 12 | TCRA_39 | TCRB_15 | Homsap TRAV8-1*01 F | Homsap TRAJ31*01 F | CAVTSNNNARLMF | Homsap TRBV7-2*01 F | | Homsap TRBJ2-1*01 F | CASSIQTSGGANEQFF |
| a40_b4 | SLA | #018 | 17 | TCRA_40 | TCRB_4 | Homsap TRAV38-2/DV8*01 F | Homsap TRAJ58*01 ORF | CAYRSALISGSRLTF | Homsap TRBV2*01 F | | Homsap TRBJ2-5*01 F | CASSEKNLAWETQYF |
| a41_b41 | SLA | #018 | 2 | TCRA_41 | TCRB_41 | Homsap TRAV38-2/DV8*01 F | Homsap TRAJ31*01 F | CAYRWNNNARLMF | Homsap TRBV7-2*01 F | | Homsap TRBJ1-4*01 F | CASSPSPSPNEKLFF |
| a43_b43 | SLA | #018 | 2 | TCRA_43 | TCRB_43 | Homsap TRAV4*01 F | Homsap TRAJ31*01 F | CLVAFNNARLMF | Homsap TRBV11-3*01 F | | Homsap TRBJ1-1*01 F | CASSQDPENTEAFF |
| a44_b41 | SLA | #018 | 5 | TCRA_44 | TCRB_41 | Homsap TRAV23/DV6*01 F | Homsap TRAJ58*01 ORF | CPGVRLTF | Homsap TRBV7-2*01 F | | Homsap TRBJ1-4*01 F | CASSPSPSPNEKLFF |
| a6_b34 | SLA | #018 | 4 | TCRA_6 | TCRB_34 | Homsap TRAV35*02 (F) | Homsap TRAJ50*01 F | CAGPETSYDKVIF | Homsap TRBV5-4*01 F | | Homsap TRBJ1-2*01 F | CASSPGQGGPRGYTF |
| a6_b49 | SLA | #018 | 18 | TCRA_6 | TCRB_49 | Homsap TRAV35*02 (F) | Homsap TRAJ50*01 F | CAGPETSYDKVIF | Homsap TRBV7-3*01 F | | Homsap TRBJ1-2*01 F | CASSSFDRGNYGYTF |
| a7_b1 | SLA | #018 | 3 | TCRA_7 | TCRB_1 | Homsap TRAV35*02 (F) | Homsap TRAJ53*01 F | CAGQDTPSGGSNYKLTF | Homsap TRBV19*01 F | | Homsap TRBJ1-1*01 F | CASRKGPEAFF |
| a8_b38 | SLA | #018 | 6 | TCRA_8 | TCRB_38 | Homsap TRAV9-2*03 (F) | Homsap TRAJ42*01 F | CAHYMNYGGSQGNLIF | Homsap TRBV7-2*01 F | | Homsap TRBJ2-5*01 F | CASSPQTGVQETQYF |
| a9 | SLA | #018 | 2 | TCRA_9 | #N/A | Homsap TRAV5*01 F | Homsap TRAJ42*01 F | CAKAGSQGNLIF | #N/A | | #N/A | #N/A |
| b11 | SLA | #018 | 2 | #N/A | TCRB_11 | #N/A | #N/A | #N/A | Homsap TRBV7-9*01 F | | Homsap TRBJ2-3*01 F | CASSGGWTVTDTQYF |
| b17 | SLA | #018 | 2 | #N/A | TCRB_17 | #N/A | #N/A | #N/A | Homsap TRBV5-1*01 F | | Homsap TRBJ2-1*01 F | CASSLALAGGAYNEQFF |
| b19 | SLA | #018 | 2 | #N/A | TCRB_19 | #N/A | #N/A | #N/A | Homsap TRBV7-2*02 F, or Homsap TRBV7-2*03 F | | Homsap TRBJ2-3*01 F | CASSLEGATDTQYF |
| b20 | SLA | #018 | 2 | #N/A | TCRB_20 | #N/A | #N/A | #N/A | Homsap TRBV5-1*01 F | | Homsap TRBJ2-4*01 F | CASSLEGENIQYF |
| b22 | SLA | #018 | 2 | #N/A | TCRB_22 | #N/A | #N/A | #N/A | Homsap TRBV7-2*01 F | | Homsap TRBJ2-7*01 F | CASSLEVLAEQYF |
| b24 | SLA | #018 | 2 | #N/A | TCRB_24 | #N/A | #N/A | #N/A | Homsap TRBV7-2*01 F | | Homsap TRBJ2-7*01 F | CASSLGGIREQYF |
| b27 | SLA | #018 | 2 | #N/A | TCRB_27 | #N/A | #N/A | #N/A | Homsap TRBV7-2*01 F | | Homsap TRBJ2-3*01 F | CASSLQAGGPDTQYF |
| b28 | SLA | #018 | 2 | #N/A | TCRB_28 | #N/A | #N/A | #N/A | Homsap TRBV7-3*01 F | | Homsap TRBJ2-3*01 F | CASSLQAGGTDTQYF |
| b3 | SLA | #018 | 4 | #N/A | TCRB_3 | #N/A | #N/A | #N/A | Homsap TRBV7-2*01 F | | Homsap TRBJ2-4*01 F | CASSDQISGHSIQYF |
| b30 | SLA | #018 | 2 | #N/A | TCRB_30 | #N/A | #N/A | #N/A | Homsap TRBV7-2*02 F, or Homsap TRBV7-2*03 F | | Homsap TRBJ2-3*01 F | CASSLQQGASDTQYF |
| b37 | SLA | #018 | 2 | #N/A | TCRB_37 | #N/A | #N/A | #N/A | Homsap TRBV7-2*01 F | | Homsap TRBJ2-1*01 F | CASSPQLAGSSNEQFF |
| b39 | SLA | #018 | 2 | #N/A | TCRB_39 | #N/A | #N/A | #N/A | Homsap TRBV7-2*01 F | | Homsap TRBJ2-3*01 F | CASSPQVGGSSTDTQYF |
| b45 | SLA | #018 | 2 | #N/A | TCRB_45 | #N/A | #N/A | #N/A | Homsap TRBV14*01 F | | Homsap TRBJ2-1*01 F | CASSQGLGTVTTEQFF |
| b51 | SLA | #018 | 2 | #N/A | TCRB_51 | #N/A | #N/A | #N/A | Homsap TRBV7-2*01 F | | Homsap TRBJ2-3*01 F | CASSSQTGATDTQYF |
| b52 | SLA | #018 | 3 | #N/A | TCRB_52 | #N/A | #N/A | #N/A | Homsap TRBV13*01 F | | Homsap TRBJ2-5*01 F | CASSSTSGGGTQYF |
| b55 | SLA | #018 | 3 | #N/A | TCRB_55 | #N/A | #N/A | #N/A | Homsap TRBV9*01 F | | Homsap TRBJ2-1*01 F | CASSVELAGYGEQFF |
| b8 | SLA | #018 | 2 | #N/A | TCRB_8 | #N/A | #N/A | #N/A | Homsap TRBV11-2*01 F | | Homsap TRBJ1-6*02 F | CASSFGQFNSPLHF |
| a4 | SLA | #004 | 3 | TCRA_4 | #N/A | Homsap TRAV5*01 F | Homsap TRAJ9*01 F | CAETAGGFKTIF | #N/A | | #N/A | #N/A |
| b17 | SLA | #004 | 2 | #N/A | TCRB_17 | #N/A | #N/A | #N/A | Homsap TRBV7-2*01 F | | Homsap TRBJ2-3*01 F | CASSLGLAGPSGTDTQYF |
| a20 | SLA | #004 | 1 | TCRA_20 | #N/A | Homsap TRAV8-3*02 (F) | Homsap TRAJ4*01 F | CAVGSGGYNKLIF | #N/A | | #N/A | #N/A |
| a33 | SLA | #004 | 1 | TCRA_33 | #N/A | Homsap TRAV8-6*02 F | Homsap TRAJ4*01 F | CAVWGSGGYNKLIF | #N/A | | #N/A | #N/A |
| b35 | SLA | #004 | 1 | #N/A | TCRB_35 | #N/A | #N/A | #N/A | Homsap TRBV7-2*01 F | | Homsap TRBJ2-3*01 F | CASSSGLAGASGTDTQYF |
| a12 | SLA | #128 | 2 | TCRA_12 | #N/A | Homsap TRAV35*02 (F) | Homsap TRAJ45*01 F | CAGQRVGGADGLTF | #N/A | | #N/A | #N/A |
| a14_b39 | SLA | #128 | 2 | TCRA_14 | TCRB_39 | Homsap TRAV9-2*02 (F) | Homsap TRAJ8*01 F | CALSDGNTGFQKLVF | Homsap TRBV3-1*01 F | | Homsap TRBJ2-3*01 F | CASSPLDTQYF |
| a27 | SLA | #128 | 2 | TCRA_27 | #N/A | Homsap TRAV8-3*02 (F) | Homsap TRAJ11*01 F | CAVENSGYSTLTF | #N/A | | #N/A | #N/A |
| a3_b62 | SLA | #028 | 14 | TCRA_3 | TCRB_62 | Homsap TRAV23/DV6*01 F | Homsap TRAJ40*01 F | CAASGPSGTYKYIF | Homsap TRBV30*01 F | | Homsap TRBJ2-4*01 F | CAWQRTGTAKNIQYF |
| a4_b23 | SLA | #028 | 2 | TCRA_4 | TCRB_23 | Homsap TRAV23/DV6*01 F | Homsap TRAJ41*01 F | CAASRSNSGYALNF | Homsap TRBV7-2*01 F | | Homsap TRBJ2-3*01 F | CASSLGLASDTQYF |
| a40_b59 | SLA | #028 | 3 | TCRA_40 | TCRB_59 | Homsap TRAV12-1*01 F | Homsap TRAJ38*01 F | CVVNNAGNNRKLIW | Homsap TRBV7-8*01 F | | Homsap TRBJ2-5*01 F | CASSVGTVQETQYF |
| a42_b10 | SLA | #028 | 5 | TCRA_42 | TCRB_10 | Homsap TRAV12-1*01 F | Homsap TRAJ40*01 F | CVVRDSGTYKYIF | Homsap TRBV5-1*01 F | | Homsap TRBJ2-3*01 F | CASSFGGAGTDTQYF |
| a7_b46 | SLA | #028 | 2 | TCRA_7 | TCRB_46 | Homsap TRAV5*01 F | Homsap TRAJ10*01 F | CAENGITGGGNKLTF | Homsap TRBV4-1*01 F | | Homsap TRBJ2-1*01 F | CASSQDPASVGSEQFF |
| a8 | SLA | #128 | 2 | TCRA_8 | #N/A | Homsap TRAV35*02 (F) | Homsap TRAJ53*01 F | CAGILSGGSNYKLTF | #N/A | | #N/A | #N/A |
| b14 | SLA | #128 | 2 | #N/A | TCRB_14 | #N/A | #N/A | #N/A | Homsap TRBV6-1*01 F | | Homsap TRBJ2-6*01 F | CASSGPYRASGANVLTF |
| b33 | SLA | #128 | 2 | #N/A | TCRB_33 | #N/A | #N/A | #N/A | Homsap TRBV11-2*03 (F) | | Homsap TRBJ2-1*01 F | CASSLVGGTTYNEQFF |
| b34 | SLA | #128 | 2 | #N/A | TCRB_34 | #N/A | #N/A | #N/A | Homsap TRBV7-9*03 F | | Homsap TRBJ1-3*01 F | CASSMTVSSGNTIYF |
| b51 | SLA | #128 | 1 | #N/A | TCRB_51 | #N/A | #N/A | #N/A | Homsap TRBV7-2*01 F | | Homsap TRBJ2-5*01 F | CASSRQGAAQETQYF |
| b53 | SLA | #128 | 2 | #N/A | TCRB_53 | #N/A | #N/A | #N/A | Homsap TRBV11-2*03 (F) | | Homsap TRBJ2-1*01 F | CASSSLQGTTVDEQFF |
| b6 | SLA | #128 | 2 | #N/A | TCRB_6 | #N/A | #N/A | #N/A | Homsap TRBV5-1*01 F | | Homsap TRBJ2-2*01 F | CASSDPSVNTGELFF |
| b66 | SLA | #128 | 2 | #N/A | TCRB_66 | #N/A | #N/A | #N/A | Homsap TRBV20-1*01 F | | Homsap TRBJ2-1*01 F | CSARVFSGGSNEQFF |
| a1_b15 | SLA | #051 | 3 | TCRA_1 | TCRB_15 | Homsap TRAV13-2*01 F | Homsap TRAJ52*01 F | CAETNAGGTSYGKLTF | Homsap TRBV20-1*02 F | | Homsap TRBJ1-1*01 F | CSASYLNTEAFF |
| a4_b5 | SLA | #051 | 2 | TCRA_4 | TCRB_5 | Homsap TRAV12-3*01 F | Homsap TRAJ57*01 F | CAMSAQGGSEKLVF | Homsap TRBV7-3*01 F | | Homsap TRBJ1-1*01 F | CASSLSGISRAFF |
| a6_b1 | SLA | #051 | 2 | TCRA_6 | TCRB_1 | Homsap TRAV12-2*01 F | Homsap TRAJ49*01 F | CAVMLPNTGNQFYF | Homsap TRBV5-1*01 F | | Homsap TRBJ2-3*01 F | CASNSLGGGDTQYF |
| a7_b8 | SLA | #051 | 7 | TCRA_7 | TCRB_8 | Homsap TRAV8-4*01 F | Homsap TRAJ15*01 F | CAVSGNQAGTALIF | Homsap TRBV4-1*01 F | | Homsap TRBJ2-5*01 F | CASSQGGTSGGLYQETQYF |
| b14 | SLA | #051 | 3 | #N/A | TCRB_14 | #N/A | #N/A | #N/A | Homsap TRBV20-1*02 F | | Homsap TRBJ2-3*01 F | CSARNAGGEDTQYF |
| a2_b6 | C.ALB | #051 | 2 | TCRA_2 | TCRB_6 | Homsap TRAV9-2*01 F | Homsap TRAJ47*02 F | CALKVEYGNKLVF | Homsap TRBV13*01 F | | Homsap TRBJ2-3*01 F | CASSLVGTDTQYF |
| a3 | C.ALB | #051 | 2 | TCRA_3 | #N/A | Homsap TRAV9-2*03 (F) | Homsap TRAJ35*01 F | CALLPPLGFGNVLHC | #N/A | | #N/A | #N/A |
| a5 | C.ALB | #051 | 2 | TCRA_5 | #N/A | Homsap TRAV25*01 F | Homsap TRAJ53*01 F | CANSGGSNYKLTF | #N/A | | #N/A | #N/A |
| b11 | C.ALB | #051 | 2 | #N/A | TCRB_11 | #N/A | #N/A | #N/A | Homsap TRBV5-1*01 F | | Homsap TRBJ2-2*01 F | CASSYDRDTGELFF |
| b12 | C.ALB | #051 | 2 | #N/A | TCRB_12 | #N/A | #N/A | #N/A | Homsap TRBV11-2*03 F | | Homsap TRBJ2-5*01 F | CASSYLGGAARGTQYF |
| b13 | C.ALB | #051 | 2 | #N/A | TCRB_13 | #N/A | #N/A | #N/A | Homsap TRBV20-1*02 F | | Homsap TRBJ1-4*01 F | CSAPKGQGTSKLFF |
| b7 | C.ALB | #051 | 3 | #N/A | TCRB_7 | #N/A | #N/A | #N/A | Homsap TRBV7-2*02 F, or Homsap TRBV7-2*03 F | | Homsap TRBJ2-3*01 F | CASSPWIQGSTDTQYF |
| a1 | C.ALB | #128 | 2 | TCRA_1 | #N/A | Homsap TRAV13-1*02 (F) | Homsap TRAJ11*01 F | CAAMVSGYSTLTF | #N/A | | #N/A | #N/A |
| a10_b8 | C.ALB | #028 | 4 | TCRA_10 | TCRB_8 | Homsap TRAV25*01 F | Homsap TRAJ38*01 F | CAGPGNAGNNRKLIW | Homsap TRBV5-1*01 F | | Homsap TRBJ2-2*01 F | CASSFARNLYDTGELFF |
| a17_b20 | C.ALB | #028 | 13 | TCRA_17 | TCRB_20 | Homsap TRAV9-2*01 F | Homsap TRAJ53*01 F | CALSGGSNYKLTF | Homsap TRBV5-1*01 F | | Homsap TRBJ2-1*01 F | CASSLEGGGAGEQFF |
| a21_b60 | C.ALB | #128 | 10 | TCRA_21 | TCRB_60 | Homsap TRAV9-2*01 F | Homsap TRAJ49*01 F | CALTLSGNQFYF | Homsap TRBV2*01 F | | Homsap TRBJ2-2*01 F | CASSVINTGELFF |
| a24 | C.ALB | #028 | 2 | TCRA_24 | #N/A | Homsap TRAV25*01 F | Homsap TRAJ42*01 F | CAPPYGGSQGNLIF | #N/A | | #N/A | #N/A |
| a36_b35 | C.ALB | #128 | 8 | TCRA_36 | TCRB_35 | Homsap TRAV8-4*01 F | Homsap TRAJ34*01 F | CAVSDRGTDKLIF | Homsap TRBV12-3*01 F | | Homsap TRBJ1-5*01 F | CASSNLGAQHF |
| a41 | C.ALB | #028 | 2 | TCRA_41 | #N/A | Homsap TRAV12-1*01 F | Homsap TRAJ38*01 F | CVVNPRAGNNRKLIW | #N/A | | #N/A | #N/A |
| b2 | C.ALB | #028 | 2 | #N/A | TCRB_2 | #N/A | #N/A | #N/A | Homsap TRBV6-5*01 F | | Homsap TRBJ2-3*01 F | CASRPGLGGDTQYF |
| b25 | C.ALB | #028 | 2 | #N/A | TCRB_25 | #N/A | #N/A | #N/A | Homsap TRBV7-2*01 F | | Homsap TRBJ2-5*01 F | CASSLLPSGGGGETQYF |
| b28 | C.ALB | #128 | 2 | #N/A | TCRB_28 | #N/A | #N/A | #N/A | Homsap TRBV18*01 F | | Homsap TRBJ1-5*01 F | CASSLQGHSNQPQHF |
| b3 | C.ALB | #028 | 8 | #N/A | TCRB_3 | #N/A | #N/A | #N/A | Homsap TRBV12-3*01 F | | Homsap TRBJ2-7*01 F | CASRPGLGIYEQYF |
| b38 | C.ALB | #128 | 2 | #N/A | TCRB_38 | #N/A | #N/A | #N/A | Homsap TRBV18*01 F | | Homsap TRBJ2-3*01 F | CASSPHSTDTQYF |
| b4 | C.ALB | #128 | 3 | #N/A | TCRB_4 | #N/A | #N/A | #N/A | Homsap TRBV12-3*01 F | | Homsap TRBJ2-5*01 F | CASRQGSETQYF |
| b43 | C.ALB | #128 | 2 | #N/A | TCRB_43 | #N/A | #N/A | #N/A | Homsap TRBV18*01 F | | Homsap TRBJ2-1*01 F | CASSPSGYNYNEQFF |
| b48 | C.ALB | #128 | 2 | #N/A | TCRB_48 | #N/A | #N/A | #N/A | Homsap TRBV4-2*01 F | | Homsap TRBJ1-2*01 F | CASSQESRGLYGYTF |
| b50 | C.ALB | #028 | 2 | #N/A | TCRB_50 | #N/A | #N/A | #N/A | Homsap TRBV12-3*01 F | | Homsap TRBJ2-2*01 F | CASSRETGTGELFF |
| b64 | C.ALB | #128 | 3 | #N/A | TCRB_64 | #N/A | #N/A | #N/A | Homsap TRBV20-1*01 F | | Homsap TRBJ1-4*01 F | CSARTRITNEKLFF |
| a14_b33 | C.ALB | #004 | 2 | TCRA_14 | TCRB_33 | Homsap TRAV17*01 F | Homsap TRAJ39*01 F | CATEFNAGNMLTF | Homsap TRBV4-2*01 F | | Homsap TRBJ2-7*01 F | CASSRASRSYEQYF |
| a2_b21 | C.ALB | #004 | 2 | TCRA_2 | TCRB_21 | Homsap TRAV29/DV5*01 F | Homsap TRAJ42*01 F | CAATGGSQGNLIF | Homsap TRBV18*01 F | | Homsap TRBJ2-1*01 F | CASSPGLAGYNEQFF |
| a29 | C.ALB | #004 | 2 | TCRA_29 | #N/A | Homsap TRAV8-4*01 F | Homsap TRAJ27*01 F | CAVSEWRAGKSTF | #N/A | | #N/A | #N/A |
| a11_b42 | C.ALB | #018 | 2 | TCRA_11 | TCRB_42 | Homsap TRAV9-2*03 (F) | Homsap TRAJ15*01 F | CALSELNQAGTALIF | Homsap TRBV4-3*01 F | | Homsap TRBJ2-7*01 F | CASSQALDSHEQYF |
| a18_b36 | C.ALB | #018 | 5 | TCRA_18 | TCRB_36 | Homsap TRAV2*01 F | Homsap TRAJ29*01 F | CAPDDGNTPLVF | Homsap TRBV18*01 F | | Homsap TRBJ2-1*01 F | CASSPPSSGGAHNEQFF |
| a19_b31 | C.ALB | #018 | 2 | TCRA_19 | TCRB_31 | Homsap TRAV19*01 F | Homsap TRAJ10*01 F | CAPRVTGGGNKLTF | Homsap TRBV7-2*01 F | | Homsap TRBJ2-7*01 F | CASSLRGAYEQYF |
| a20_b58 | C.ALB | #018 | 6 | TCRA_20 | TCRB_58 | Homsap TRAV17*01 F | Homsap TRAJ30*01 F | CATQRQGDKIIF | Homsap TRBV9*01 F | | Homsap TRBJ2-1*01 F | CASSVLPGGGAYNEQFF |
| a21_b32 | C.ALB | #018 | 6 | TCRA_21 | TCRB_32 | Homsap TRAV17*01 F | Homsap TRAJ37*01 F | CATRSGNTGKLIF | Homsap TRBV7-2*01 F | | Homsap TRBJ1-4*01 F | CASSLSQGGAEKLFF |
| a22 | C.ALB | #018 | 2 | TCRA_22 | #N/A | Homsap TRAV39*01 F | Homsap TRAJ30*01 F | CAVDTSRRDDKIIF | #N/A | | #N/A | #N/A |
| a25_b13 | C.ALB | #018 | 14 | TCRA_25 | TCRB_13 | Homsap TRAV20*02 (F) | Homsap TRAJ49*01 F | CAVFNTGNQFYF | Homsap TRBV7-6*01 F | | Homsap TRBJ2-1*01 F | CASSHRSGRQFF |
| a3_b40 | C.ALB | #018 | 4 | TCRA_3 | TCRB_40 | Homsap TRAV13-2*01 F | Homsap TRAJ52*01 F | CAEMGGTSYGKLTF | Homsap TRBV11-2*01 F | | Homsap TRBJ2-6*01 F | CASSPQVSGANVLTF |
| a33_b26 | C.ALB | #018 | 11 | TCRA_33 | TCRB_26 | Homsap TRAV41*01 F | Homsap TRAJ39*01 F | CAVRRNNAGNMLTF | Homsap TRBV5-5*02 (F) | | Homsap TRBJ2-7*01 F | CASSLLGAYEQYF |
| a33_b31 | C.ALB | #018 | 6 | TCRA_33 | TCRB_31 | Homsap TRAV41*01 F | Homsap TRAJ39*01 F | CAVRRNNAGNMLTF | Homsap TRBV12-3*01 F | | Homsap TRBJ2-7*01 F | CASSLRGAYEQYF |
| a33_b47 | C.ALB | #018 | 6 | TCRA_33 | TCRB_47 | Homsap TRAV41*01 F | Homsap TRAJ39*01 F | CAVRRNNAGNMLTF | Homsap TRBV4-2*01 F | | Homsap TRBJ2-7*01 F | CASSQVGAYEQYF |
| a34_b48 | C.ALB | #018 | 8 | TCRA_34 | TCRB_48 | Homsap TRAV41*01 F | Homsap TRAJ39*01 F | CAVRTNNAGNMLTF | Homsap TRBV12-3*01 F | | Homsap TRBJ2-7*01 F | CASSRAKGAYEQYF |
| a36_b18 | C.ALB | #018 | 3 | TCRA_36 | TCRB_18 | Homsap TRAV8-4*03 (F) | Homsap TRAJ34*01 F | CAVSGGNTDKLIF | Homsap TRBV12-3*01 F | | Homsap TRBJ2-3*01 F | CASSLAPGAGTQYF |
| a37_b46 | C.ALB | #018 | 5 | TCRA_37 | TCRB_46 | Homsap TRAV12-2*02 (F) | Homsap TRAJ41*01 F | CAVSNSNSGYALNF | Homsap TRBV4-2*01 F | | Homsap TRBJ2-1*01 F | CASSQRIRTGPSSYNEQFF |
| a4 | C.ALB | #018 | 2 | TCRA_4 | #N/A | Homsap TRAV24*01 F | Homsap TRAJ49*01 F | CAFGNTGNQFYF | #N/A | | #N/A | #N/A |
| a42_b36 | C.ALB | #018 | 9 | TCRA_42 | TCRB_36 | Homsap TRAV26-1*01 F | Homsap TRAJ38*01 F | CIVRNAGNNRKLIW | Homsap TRBV18*01 F | | Homsap TRBJ2-1*01 F | CASSPPSSGGAHNEQFF |
| a45 | C.ALB | #018 | 4 | TCRA_45 | #N/A | Homsap TRAV8-2*01 F | Homsap TRAJ39*01 F | CVVRGPNAGNMLTF | #N/A | | #N/A | #N/A |
| a5_b61 | C.ALB | #018 | 5 | TCRA_5 | TCRB_61 | Homsap TRAV25*01 F | Homsap TRAJ32*02 F | CAGGGGYGGATNKLIF | Homsap TRBV20-1*01 F | | Homsap TRBJ1-2*01 F | CSARTYQRKTNYGYTF |
| b12 | C.ALB | #018 | 2 | #N/A | TCRB_12 | #N/A | #N/A | #N/A | Homsap TRBV2*01 F | | Homsap TRBJ1-3*01 F | CASSGSGGNTIYF |
| b16 | C.ALB | #018 | 5 | #N/A | TCRB_16 | #N/A | #N/A | #N/A | Homsap TRBV5-5*02 (F) | | Homsap TRBJ2-7*01 F | CASSLAGAYEQYF |
| b2 | C.ALB | #018 | 3 | #N/A | TCRB_2 | #N/A | #N/A | #N/A | Homsap TRBV6-1*01 F | | Homsap TRBJ1-2*01 F | CASRVSPRGYTF |
| b25 | C.ALB | #018 | 2 | #N/A | TCRB_25 | #N/A | #N/A | #N/A | Homsap TRBV7-9*01 F | | Homsap TRBJ2-4*01 F | CASSLGGVVAKNIQYF |
| b50 | C.ALB | #018 | 2 | #N/A | TCRB_50 | #N/A | #N/A | #N/A | Homsap TRBV12-3*01 F | | Homsap TRBJ2-5*01 F | CASSSIGGETQYF |
| b56 | C.ALB | #018 | 3 | #N/A | TCRB_56 | #N/A | #N/A | #N/A | Homsap TRBV2*01 F | | Homsap TRBJ1-1*01 F | CASSVFGSMNTEAFF |
| b59 | C.ALB | #018 | 2 | #N/A | TCRB_59 | #N/A | #N/A | #N/A | Homsap TRBV4-1*01 F | | Homsap TRBJ1-6*02 F | CASTQSTGNNSPLHF |

**Extended Table 3. Sequence of SLA- and C.Alb-specific TCRab from identified clones**
